# Inference with selection, varying population size and evolving population structure: Application of ABC to a forward-backward coalescent process with interactions

**DOI:** 10.1101/819318

**Authors:** Clotilde Lepers, Sylvain Billiard, Matthieu Porte, Sylvie Méléard, Viet Chi Tran

## Abstract

Genetic data are often used to infer history, demographic changes or detect genes under selection. Inferential methods are commonly based on models making various strong assumptions: demography and population structures are supposed *a priori* known, the evolution of the genetic composition of a population does not affect demography nor population structure, and there is no selection nor interaction between and within genetic strains. In this paper, we present a stochastic birth-death model with competitive interaction to describe an asexual population, and we develop an inferential procedure for ecological, demographic and genetic parameters. We first show how genetic diversity and genealogies are related to birth and death rates, and to how individuals compete within and between strains. This leads us to propose an original model of phylogenies, with trait structure and interactions, that allows multiple merging. Second, we develop an Approximate Bayesian Computation framework to use our model for analyzing genetic data. We apply our procedure to simulated and real data. We show that the procedure give accurate estimate of the parameters of the model. We finally carry an illustration on real data and analyze the genetic diversity of microsatellites on Y-chromosomes sampled from Central Asia populations in order to test whether different social organizations show significantly different fertility.

## 1 Introduction

Genetic polymorphism within and between taxa is commonly used for estimating population structures Goldstein and Chikhi (2002); Müller *et al.* (2017) or demographic changes Beichman *et al.* (2018), to infer population history, migration patterns, or to search for genes under selection Stephan (2016). These methods are mostly based either on the site frequency spectrum, the identity per state or descent, or on summary statistics in an Approximate Bayesian Computation (ABC) framework Beaumont *et al.* (2002). Statistical testing and model selection are generally performed under simplifying assumptions which allows computations of quantities such as the likelihood of a model, in particular under neutrality. For instance, demography generally follows a Wright-Fisher model or a birth-death model without interactions.

Under the Wright-Fisher model, the population size is supposed deterministic: it is known at any given time and independent of the composition of the population, *i.e.* it is supposed that the mechanisms underlying the population size variations are extrinsic and without noise. Individuals thus compete for space but the carrying capacity of the environment does not change because of the evolution of the population itself, or because of extrinsic or intrinsic stochasticity. In birth-death models, population size can vary but populations can grow indefinitely because individuals do not interact. In addition, the Wright-Fisher and birth-death models are supposed neutral *i.e.* the reproduction and survival rates do not depend on the genetic lineage (but see a recent birth-death model without interactions where rates can depend on mutations Rasmussen and Stadler (2019)).

Yet, the assumptions of neutrality, extrinsic control of population size or non-interacting individuals are certainly often violated. For instance, genealogies of the seasonal influenza virus show important departure from neutrality which might suggest that selection and interaction between lineages are important enough to significantly affect evolution and the shapes of the phylogenetic trees Bedford *et al.* (2011); Strelkowa and Lässing (2012). Reproduction rates and carrying capacity have also been shown to depend on strains in the domesticated yeasts Spor *et al.* (2009). In addition, the ecological literature contains many cases where competitive interactions vary among strains or species Gallieni (2017). Developing models and inference methods which relax such hypotheses is thus a contemporaneous challenge, in order to improve our knowledge of the history and ecological features of species and populations. As emphasized by Frost *et al.* (2015), this challenge is in particular important for the analysis of phylodynamics in clonal species such as viruses.

Some of these assumptions have been already relaxed. For instance, Rasmussen and Stadler (2019) developed a model where reproductive and death rates can differ between lineages which can emerge because of spontaneous mutations. They applied their method on Ebola and influenza viruses in order to have estimate of fitness effects of mutations from phylodynamics. Indeed, variation of death and birth rates between lineages can affect viruses phylogenies, which can be detected and used to infer the effect of mutations. However, they supposed no interaction between lineages, discarding a possible effect of competition between viruses strains.

In this paper, we present a model and an inference method which allow the relaxation of several of these assumptions. First, in Sections 2.1 and 2.2, we develop a stochastic process describing the eco-evolution of a structured population with ecological feedbacks (based on Billiard *et al.* (2015)). This model takes into account i) mutations that can affect birth, death and competitive rates; ii) explicit competitive interactions between lineages; iii) population size depending on the genetic composition of the population, *i.e.* the carrying capacity depends on the ecological properties of existing strains (their birth, death and competitive rates); iv) the structure of the population is not fixed and can change depending on the genetic evolution of the population. The model assumes reproduction is asexual, that mutations affecting fitness are rare, and that neutral mutation follows an intermediate timescale between reproduction and death rates (the ecological timescale) and the rate at which mutations affecting fitness appear (the evolutionary timescale). Second, in Section 2.3, a new forward-backward coalescent process is proposed to describe the phylogenies in such a population. The forward step accounts for interactions, demography and evolution of trait structures, defining the skeleton on which the phylogenies of sampled individuals can be reconstructed in the backward step. Such a forward-backward approach has been used in the case of nested coalescent models, *e.g.* Benítez *et al.* (2018); Duchamps (2018); Verdu *et al*. (2009), but in our case interactions within and between lineages, ecological feedbacks between selection and population size, and multiple merging are taken into account. Contrarily to Λ-coalescent models proposed in the literature Donnelly and Kurtz (1999b); Pitman (1999); Sagitov (1999), multiple merging here are not due to high reproduction variances but they appear as a consequence of natural selection via mutation-competition and timescales. Third, in Section 3, we develop an ABC framework in order to estimate the parameters of the model from genetic diversity data. We show how ecological parameters such as individual birth and death rates, and competitive abilities can be estimated. Finally, we reanalyze genetic data from Y-chromosomes sampled in Central Asia human populations (Chaix *et al*. (2007); Heyer *et al.* (2015)) in order to test whether different social organizations can be associated with difference in fertility.

## 2 The forward-backward coalescent model

### 2.1 A stochastic birth-death model with competition

We assume a population of clonal individuals, characterized, on the one hand, by a trait, *i.e.* a vector of genetically determined variables *x* ∈ 𝒳 ⊂ ℝ^*d*^, which affects the demographic processes such as birth, death and competitive interactions between individuals, and, on the other hand, by a vector of genetic markers *u* ∈ 𝒰 ⊂ ℝ^*q*^, supposed neutral (*i.e. u* does not affect demographic processes). 𝒳 and 𝒰 respectively represent the sets of possible values of the trait and the neutral markers. Trait *x* can represent a proxy for strains, lineages, species or subpopulations. We assume that the marker *u* is totally linked to the trait *x* (no recombination). The number of individuals in the population at time *t* is denoted 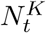, where *K* is a scaling parameter which controls the relationship between the demographic and genetic parameters and variables. For instance, regarding competition, one can consider that each individual has a weight 1*/K* such that when *K* increases, competition between individuals decreases and the population size increases.

The evolution of the population is a stochastic process in continuous time which depends on the rates of all possible demographic and genetic processes. Individuals with trait *x* give birth at rate *b*(*x*) and die at rate 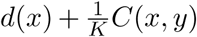, where *d*(*x*) is the intrinsic death rate, and 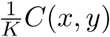 is the additional death rate due to the competitive effect of a single individual with trait *y*. At birth, offspring’ traits and markers can change by mutation with probability *p*_*K*_ and *q*_*K*_, respectively. The trait and marker mutation rates are respectively supposed such that *p*_*K*_ = 1*/K*^2^ and

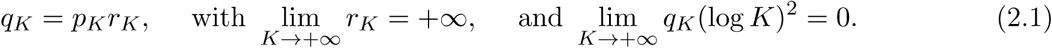

In a few words, the previous assumptions mean that ecological processes occur faster than mutations, and that mutations on the neutral marker *u* occur faster than mutations on the trait *x*. The model thus suppose three timescales (more details are given below). After mutation on the trait *x*, the offspring’s trait is *x* + *h* with *h* randomly drawn in a distribution *m*(*x, h*)*dh*. The effect of mutations on the neutral marker are supposed to follow a Gaussian distribution with mean 0 and variance 1*/K* (but alternatives are possible, see Billiard *et al.* (2015)). Notice that we do not need to assume small mutations on the trait *x, i.e.* selection can be strong.

### 2.2 Genetics diversity in an eco-evolutionary dynamics with three timescales

We construct from the microscopic model of Section 2.1 a precise tractable approximation built on the separations of timescales. Under our assumptions about how fast ecological and genetic processes occur, the population evolves following three timescales. First, the ecological timescale: birth and death rates occur at a rate of order *K, i.e.* the size of the population. Second, marker mutations arise slightly slower than the ecological timescale: they occur at a rate of order *Kq*_*K*_ = *r*_*K*_*/K* = *o*(*K/*(log *K*)^2^) (Eq. 2.1). Finally, mutations on the trait under selection occur at the slowest timescale, at a rate of order 1*/K* = *o*(*r*_*K*_*/K*). This reflects for instance that a large proportion of a genome is not composed of traits under selection. Such an assumption is necessary in the case of evolving clonal populations otherwise neutral diversity would be lost at every adaptive step. For instance, despite a very rapid evolution and adaptation, the influenza virus shows a large diversity within season Neher and Bedford (2015).

When the population is large, *i.e. K* → *∞*, the evolution of the population can be decomposed into the succession of invasions of favorable mutations on the trait *x*, because ecological processes are very fast, and the population jumps from one state to another. This corresponds to the Adaptive Dynamics framework described in Billiard *et al.* (2015) following the pioneer works of Metz *et al.* (1996); Champagnat (2006); Champagnat and Méléard (2007). Rescaling time by a factor *K* is similar to the rescaling commonly chosen under the Kingman’s coalescent Kingman (1982). The neutral marker also evolves between each adaptive jump, at a faster timescale that is compensated by mutations of small effect (of order 1*/K*). Since the ecological parameters change at every adaptive jump on the trait *x* (the birth rate, death rate and the population size change), the evolution of the neutral marker also changes. Hence, even if the marker is neutral, its own evolution depends on the state of the population at a given time, especially on the competitive interactions between individuals *C*(*x, y*). Overall, the joint eco-evolutionary dynamics of the neutral marker and the selected traits can be approximated by the so-called Substitution Flewing-Viot Process (SFVP, see Billiard *et al.* (2015)). The properties of the SFVP approximation are described in the following. Mathematical details and proofs are given in Appendix A.

#### Distribution of the trait x between two adaptive jumps

At the ecological timescale, when the population is large (*K* → *∞*), *p* strains with traits *x*_1_, … *x*_*p*_ can coexist. Between two adaptive jumps, the trait distribution in the population remains almost constant. Indeed, the size of subpopulations can vary but are expected to stay close to their equilibria 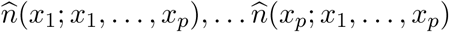, given by the following competitive Lotka-Volterra system of ordinary differential equations (ODE) that approximates the evolution in the ecological timescale:

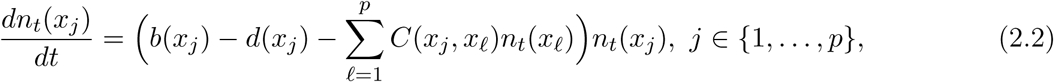

where 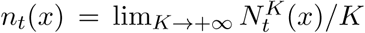 is the renormalized population size of strain with trait *x* with *K* → *∞. n*_*t*_ can be seen as the density of individuals, which is the continuous limit of the population size when it is large and composed of infinitesimal individuals. To ensure that the ODE system in Eq. 2.2 admits uniquely defined equilibria (thus excluding cyclical or chaotic dynamics), we must assume that competitive interactions are such that the matrix (*C*(*x*_*i*_, *x*_*j*_))_1≤ *i,j*≤ *p*_ is symmetric positive definite. Notice that the equilibrium 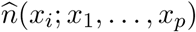 of the population of the strain with trait *x*_i_ depends on the whole trait structure of the population which is in turn defined entirely by the set of traits present in the population (the arguments of 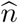 given after the semicolon).

#### Change of the distribution of the trait x during an adaptive jump

At time *t*, in the timescale of mutation rates on the trait *x*, the population is composed of *p* strains with traits *x*_1_, … *x*_*p*_ and respective sizes 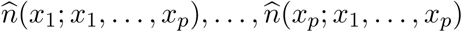. An independent exponential random clock with parameter 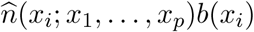 is ‘attached’ to each strain *i*. Populations remain close to their equilibria until the first clock rings. If strain *i* clock rings first, then a mutation on the trait occurs and a new strain is introduced in the population with trait *x*_*i*_ + *h* where *h* is drawn in the distribution *m*(*x*_*i*_, *h*)*dh*. Whether the mutant strain invades or not the population depends on its invasion fitness defined by

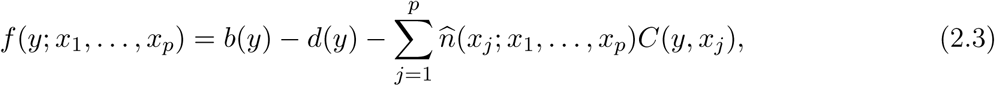

which gives the probability of invasion

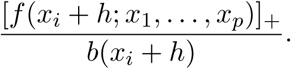

In case the mutant strain invades, the population jumps to a new state given by the solution of the Lotka-Volterra ODE system (Eq. 2.2) updated with the introduction of the mutant strain 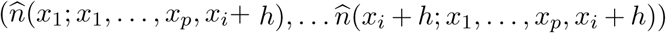 In the new equilibrium, some former traits *x*, …, *x*_*p*_ may be lost. The evolution of the trait can thus be described by the succession of the adaptive jumps of the population from one state to another, which is called Polymorphic Evolutionary Sequence (PES, Champagnat and Méléard (2007)).

#### Evolution of the neutral marker

The marker distribution within each strain evolves as a so-called Fleming-Viot process. Since we assume that the marker mutates at a faster timescale than the trait, and that mutations are small, the Fleming-Viot process describing the evolution of the neutral marker can be approximated by a diffusive limit. Here, we assume that the effect of mutations on the marker are drawn in a Gaussian distribution. Consequently, the diffusive limit distribution of the marker is given by the Fleming-Viot superprocess (Dawson (1993); Dawson and Hochberg (1982); Donnelly and Kurtz (1999a, 1996); Etheridge (2000) and App.A.4). If the mutations on the marker were discrete and take a fixed number of possible values, the diffusive limit of the marker distribution would be given by a Wright-Fisher diffusion (e.g. Crow and Kimura (1970); Nagylaki (1989)).

When the mutant strain with trait *x*_*i*_ + *h* invades the population, say at time 0, an adaptive jump occurs. Let us denote by *u* the marker of the first mutant individual (*x*_*i*_ + *h, u*). The initial condition of the Fleming-Viot *F*_*t*_(*x*_*i*_ + *h*,.) describing the distribution of the neutral marker within strain *i* and trait *x* = *x*_*i*_ + *h*, is composed of one individual with marker *u*. The evolution of the marker distribution within this strain is given by the probability measure 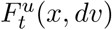 on 𝒰 at time *t\**. In other words, 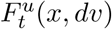 is the distribution at time *t* of the marker values within the strain with trait *x* = *x*_*i*_ + *h* given the initial value *u*. This distribution changes with time depending on the supposed mutation kernel on the marker, on the birth and death rates of individuals with trait *x*_*i*_ + *h*, and on the competitive interactions *C*(*x*_*i*_ + *h, y*) with all the other individuals of any trait value *y ∈ {x*_1_, …, *x*_*p*_, *x*_*i*_ + *h}*. Between two adaptive jumps, how the distribution 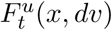 evolves with time is given by the following stochastic differential equation (Billiard *et al.* (2015))

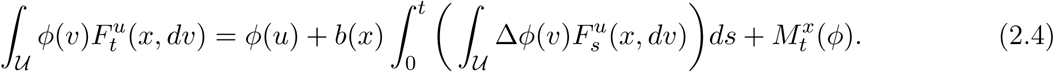

The left side of the equation can be seen as the expectation of the distribution of the marker value at time *t*, where *ϕ* is a test function (supposed twice differentiable on 𝒰). The right side of the equation tells what is the expected form of the distribution. The first term on the right side gives the initial conditions: the first mutant with trait *x*_*i*_ + *h* has a marker value *u*, hence the initial condition for the distribution is *ϕ*(*u*). The second term on the right side integrates the changes of the distribution which are only due to mutations on the marker between time 0 (the invasion time of *x* = *x*_*i*_ + *h*) and *t*. Since mutation only occurs at birth, the rate at which *F* changes with mutation is proportional to the birth rate *b*(*x*) = *b*(*x*_*i*_ + *h*). Within the integral, Δ*ϕ*(*v*) is the Laplacian of the function *ϕ* which gives the rate of change of *F* in all the dimensions of the marker values (which depends on the assumptions made on the mutation kernel). The last term 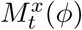 on the right side gives the changes of *F* which are due to the ecological processes, *i.e.* the fluctuations due to the birth and death of the individuals with trait *x* = *x*_*i*_ + *h*. 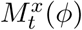 is a martingale *i.e.* a square integrable random variable with mean 0 and variance

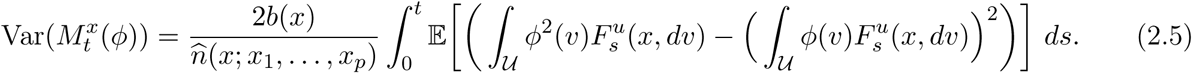

The fraction in the right hand side (r.h.s.) of Eq. 2.5 corresponds to the demographic al variance 2*b*(*x*) divided by the effective population size

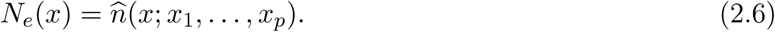

The population effective size, which partially governs the evolution of the diversity at the neutral marker, depends on the trait value *x*, but also on the whole trait distribution *x*_1_, …, *x*_*p*_, *x*_*i*_ + *h*, given by the solution of the Lotka-Volterra ODE system (Eq. 2.2). In particular, it means that the variance in the neutral diversity within the strain with trait *x* depends on the competitive interactions of the latter with all the other strains. Eq. 2.4 is complex because it is a very general form that can be applied in many different situations regarding the genetic and ecological processes.

### 2.3 Genealogies in a forward-backward coalescent with competitive interactions

In the previous section, we described how ecological processes and genetic diversity are linked in our model. We now go a step further by showing how the genealogies of a sample of size *n* individuals are determined in our model. Adaptive jumps of the PES change how the population is structured. Either the resident trait is replaced by an invading mutant, or several subpopulations are stably maintained at equilibrium and the population is polymorphic for the trait. Each time the trait evolves, the birth and death rates and the competitive interactions change, as well as the population and subpopulation sizes. Since the evolution of the trait depends on the current distribution of the traits and their respective population sizes, the PES tree should be constructed forward in time. The PES thus yields a genealogical tree based on the traits over time. Consider a subpopulation *i* with its own trait *x*_*i*_ that is introduced and maintained during the PES. This subpopulation has its own coalescent rate on the markers which depends on its own reproductive rate *b*(*x*_*i*_) and its equilibrium population size *N*_*e*_(*x*_*i*_), as shown by Eq. 2.6, that depends on the other traits. Hence the coalescent rate within a population also depends on the distribution of the trait in the whole population. Genealogies are thus expected to be different among the different strains and between different adaptive jumps of the PES. Between these adaptive jumps however, the population structures on the trait and the population size are supposed fixed. Hence, on these time intervals and given the distribution of the trait in the population, the genealogies within each strain can be constructed backward.

In our model, genealogies are thus piece wise-defined and constructed by dividing time between intervals separating adaptive jumps of the PES. As shown earlier, the marker distribution on such intervals is a Fleming-Viot process. Hence the within-strain genealogies follow a Kingman coalescent processes with a rate depending on *N*_*e*_(*x*) (EQ. 2.6), and such that at the time of an adaptive jump, multiple coalescence occur since a single individual gives birth to a new strain. This creates a bottleneck for the lineages with a hitchhiking phenomenon for the marker carried by the successful mutant. The evolution of the population can thus finally be described by a forward-backward coalescent process more precisely defined as follows.

Forward step First, a PES in forward time is considered where the successive adaptive jump times are denoted by (*T*_*k*_)_*k∈{*1,*…J}*_, with *T*_0_ = 0 and *J* is the number of jumps that occurred before time *t*.

Backward step Second, given the PES during the time interval [*T*_*k*_, *T*_*k*+1_) (*k ∈ {*0, … *J* − 1*}*), the genealogy of the sample of *n* individuals drawn independently in the population at time *t* is constructed backward in time. Assuming that the population is composed of the traits *{x*_1_, … *x*_*p*_*}* on this time interval, the genealogy of the individuals within the strain with trait *x*_*i*_ is obtained by simulating a Kingman coalescent with coalescence rate 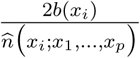 (Eq. 2.5). In absence of interaction *C*(*x, y* = 0), and with non-varying coalescence rates, the resulting process is the nested Kingman coalescent (e.g. Benítez *et al.* (2018); Duchamps (2018); Lambert and Schertzer (2018); Meizis (2018)). At time *T*_*k*_, all lineages in the subpopulation of strain with trait *x*_*i*_, that is created at this time, instantaneously coalesce because a single mutant is always at the origin of a new strain during the PES. Note that coalescence is instantaneous under the assumptions underlying the PES, *i.e.* at the timescale governing the evolution of the trait, the transition to fixation of the mutant trait is negligible and all the lineages stemming from this period seem to coalesce instantaneously. The allelic state at the marker can finally be determined given the previously constructed genealogy, depending on the mutational model considered.

A more formal definition of the coalescent and associated proofs are given in App. A.3. A simulation algorithm for the construction of genealogies under our model is given in App. A.4.

## 3 ABC inference in an eco-evolutionary framework

We showed in the previous sections that the genetic structure of a sample of *n* individuals can be related to the parameters of our eco-evolutionary model. We now aim at using this framework to infer genealogies, ecological and genetic parameters from genetic and/or phenotypic data sampled in a population at time *t*. In other words, given a dataset containing the genotype at the marker *u* and the genotype or phenotype at the trait *x* for the *n* sampled individuals, we want to infer the parameters of the model: birth, death and competitive interaction rates, mutation rates, etc. Since we have only a partial information of the population (*n* individuals are sampled and possible extinct lineages might be unobserved), the likelihood of a model given the data has no explicit form. Given a possible genealogy of the *n* individuals, an infinite number of continuous genealogical trees could be obtained from the model. The likelihood of each tree depends on the number and the traits of the different subpopulations during the history of the population, including the unobserved and extinct ones. Because summing over all possible unobserved data (number of unobserved and extinct lineages with their traits and adaptive jump times) is not feasible in practice, we have to make inference without likelihood computations.

An alternative to likelihood-based inference methods is given by the Approximate Bayesian Computation (ABC) (Beaumont *et al.* (2002, 2009)), which relies on repeated simulations of the forward-backward coalescent trees (Section 2.3). In the following, we briefly give a general presentation of the application of the ABC method to our model. We then apply the method to simulations of a toy model (the Dieckmann-Doebeli model) and to real data (genetic data on microsatellites on the Y chromosomes of populations from Central Asia populations, with their social and geographic structures).

### 3.1 ABC estimation of the ecological parameters based on the genealogical tree

The dataset denoted **z** contains the genotype and/or phenotype on the trait *x* and the marker *u* for each of the *n* sampled individuals. The trait *x* can be geographic locations, species or strain identity, size, color, genotypes or anything that affect the ecological parameters and fitness. The marker *u* can also be genotypic or phenotypic measures, discrete or continuous, qualitative or quantitative, but with no effect on fitness (the marker is supposed neutral). Our goal is to use the dataset **z** to estimate the parameters of the model denoted *θ* (in our case, birth and death rate, competition kernel, mutation probabilities and kernel) using an ABC approach. To do so, the following procedure is repeated a large number of times:

1^*st*^ step. A parameter set *θ*_*i*_ is drawn in a prior distribution *π*(*dθ*);

2^*st*^ step. A PES and its neutral nested genealogies of the *n* sampled individuals are simulated in each model associated with the parameters *θ*_*i*_;

3^*rd*^ step. A set of summary statistics *S*_*i*_ is computed from the data simulated under *θ*_*i*_, for each *i*.

The posterior distribution of the model is then approximated by comparing, for each simulation *i*, the simulated summary statistics *S*_*i*_ to the ones from the real dataset and by computing for each parameter *θ*_*i*_ a weight *W*_*i*_ that defines the approximated posterior distribution (see Formula B.1 in Appendix). Three categories of summary statistics have been used, each associated with a different aspect of the genealogical tree (the complete list of summary statistics is given in the Appendix D):

- The trait distribution describing the strains diversity and their abundances (*e.g.* number of strains, the mean and variance of strains abundance, …);
- The marker distribution in the sampled population describing the neutral diversity within each sampled strain (*e.g.* the M-index, *F*_*st*_, Nei genetic distances,…);
- The shape of the genealogy (*e.g.* most recent common ancestor, length of external branches, number of cherries, …).

Depending on the dataset and the information that is available for a given population, four scenarios can be encountered:

#### Scenario 1. Complete information

The evolutionary history of the trait and the genealogies, populations and subpopulations abundances, values of the sampled individuals on the trait *x* and the marker *u*. This situation certainly never occurs but it is a reference which allows to evaluate the expected ABC estimation in a perfect situation where all information is available. This situation can also include cases where independent information can be added such as fossil records;

#### Scenario 2. Population information

Total population abundance, values of the trait *x* and marker *u* of the sampled individuals. The estimations given with those statistics represent the estimations one could expect with a complete knowledge of the present population;

#### Scenario 3. Sample information

The number of sampled sub-populations, the values of the trait *x* and the marker *u* of the sampled individuals;

#### Scenario 4. Partial sample information

Only the number of sampled sub-populations and the values of the marker *u* of the sampled individuals.

The four situations will be compared regarding the quality of the ABC estimations of the model parameters.

### 3.2 Application 1: Inference of the parameters in the Dieckmann-Doebeli model

In this section, we applied the ABC statistical procedure on the traits distribution and their phylogenies generated by a simple eco-evolutionary model Roughgarden (1979); Dieckmann and Doebeli (1999); Champagnat *et al.* (2006). The birth rate of an individual with trait *x* is 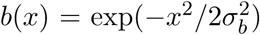, the individual natural death rate is constant *d*(*x*) = *d*_*C*_, and the competition between two individuals with traits *x* and *y* is 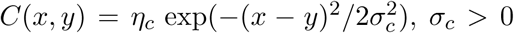, *σ*_*c*_ > 0. The trait and marker spaces are assumed bounded such as 𝒳 = [−1, 1] and 𝒰 = [−2, 2], respectively. The effect of a mutation on the trait *x* is randomly drawn in a Gaussian mutation kernel with mean 0 and variance 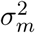 (values outside 𝒳 are excluded). The effect of mutations on the marker *u* is drawn in a centered Gaussian distribution with variance 1*/K*. The distribution of the phylogenies depends on the parameter *θ* = (*p, q, σ*_*b*_, *σ*_*c*_, *σ*_*m*_, *d*_*c*_, *η*_*c*_, *t*_*sim*_), where *t*_*sim*_ is the duration of the PES (*t*_*sim*_ is not known *a priori* and must be considered as a nuisance parameter).

#### 3.2.1 Posterior distribution and parameters estimation

We ran *N* = 400 000 simulations with identical prior distributions and scaling parameter *K* = 1000 (see details in App. B). Chosen parameter sets and prior distributions are given in Appendix A.4. We randomly chose four simulations runs among the *N* simulations as *pseudo-data* sets (Fig. 3.1). All other simulations runs were used for the parameters estimates. Fig 3.2 shows the posterior distribution for one of the the pseudo-dataset (see App. D.5 for full results). Our results show that estimates based on all statistics (Scenario 1, blue distribution) are not always the most accurate, suggesting that some of the descriptive statistics introduce noise and worsen estimate accuracy. On the contrary, knowledge about how population is trait-structured importantly improves estimation (compare orange *vs.* red posterior distributions).

**Figure 3.1:**
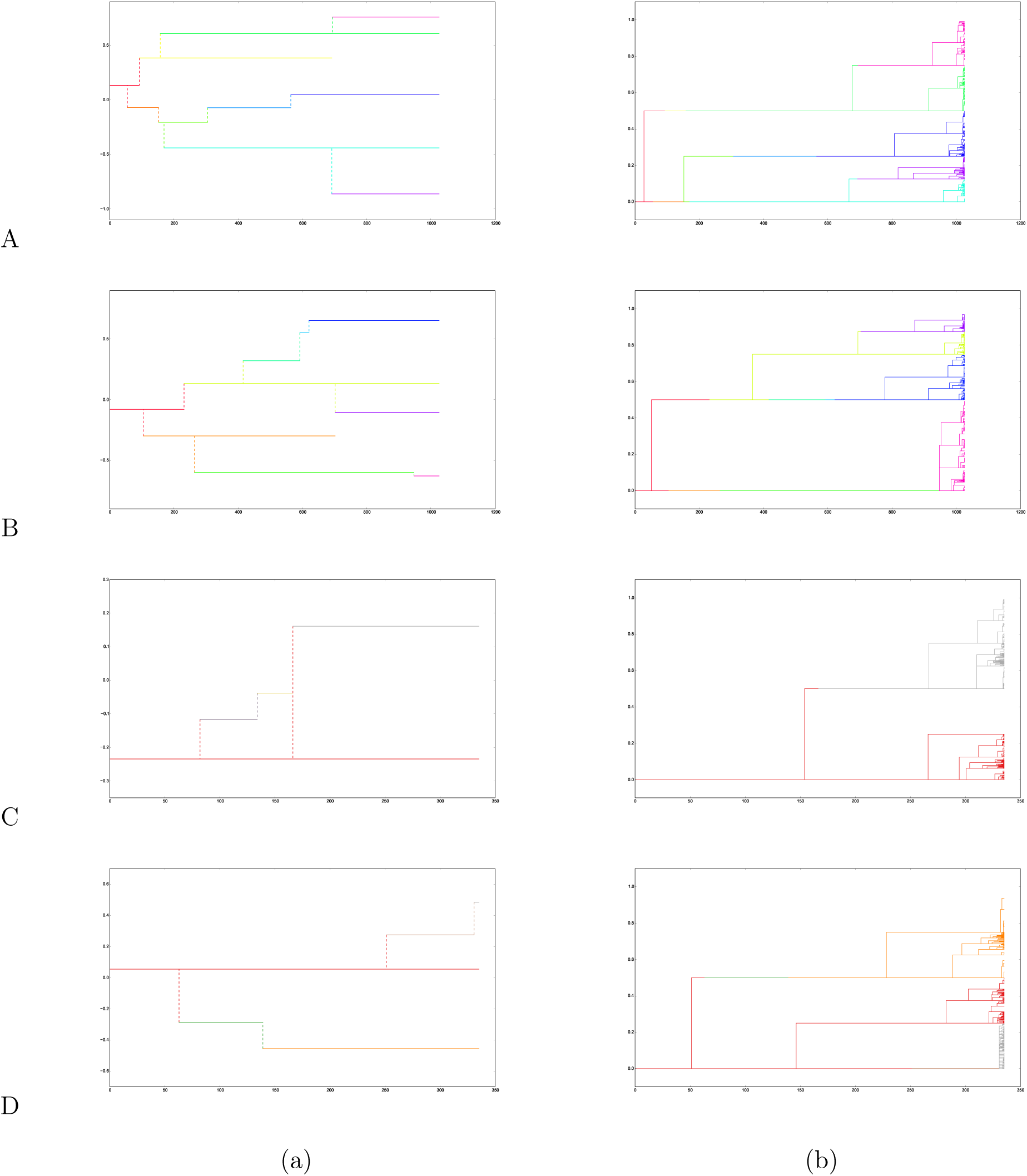
*Dynamics of (a) the trait x and (b) the neutral marker u of the four pseudo-data sets randomly sampled among N* = 400, 000 *simulations runs of the Doebeli-Dieckmann’s model. Figures show the Substitution Fleming-Viot Process (SFVP) and the nested phylogenetic tree of n individuals sampled at the final time of the simulation (Parameter sets are given in App. Table 1)*. *(a): The trait x follows a Polymorphic Evolutionary Substitution (PES) process introduced in Champagnat and Méléard (2007). (b): The genealogies of the marker u follow a forward-backward coalescent process nested in the PES tree as described in Section 2.3. The colors refer to the lineage to which one individual belong shown in (a).*

**Figure 3.2:**
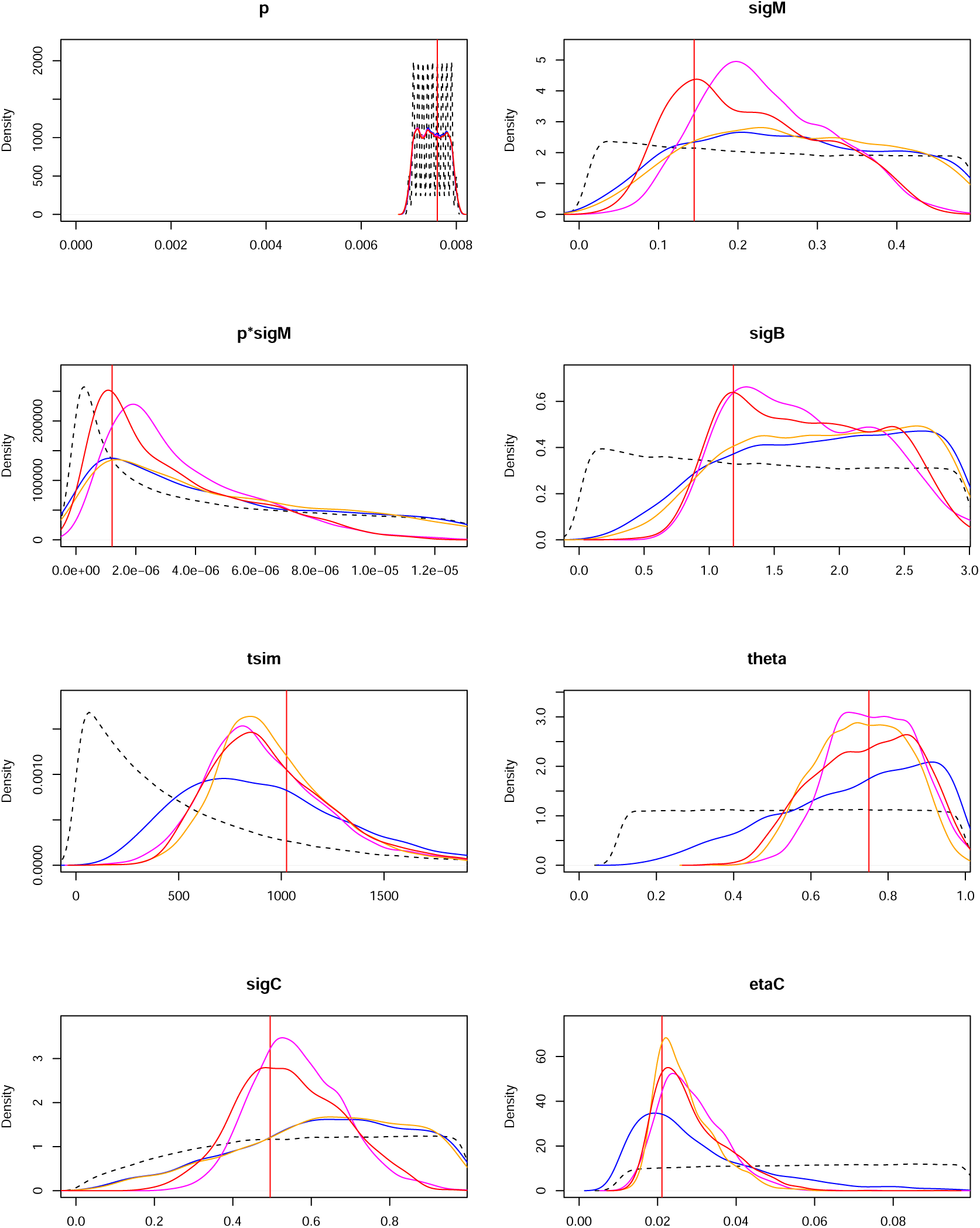
Prior and posterior distributions (pseudo-data set A in Fig. 3.1). Black dashed curve: prior distribution; Vertical red line: true value. The different colors correspond to different scenario regarding which data are available or not: Blue, Scenario 1 (All descriptive statistics are available); Pink, Scenario 2 (data from the totality of the population); Red, Scenario 3 (data from a sample of the population); Orange, Scenario 4 (data from a sample of the population, the traits x is not known). Results for other pseudo-data sets are given in App.D.5

#### 3.2.2 Do coalescent trees significantly differ from a Kingman’s coalescent?

The genealogies generated by the forward-backward coalescent under a Doebeli-Dieckmann’s model are expected to differ from a standard or renormalized Kingman’s coalescent for various reasons: i) there are multiple instantaneous coalescence events when a new lineage appears, ii) coalescence rates differ among lineages, creating asymmetries in the phylogenetic tree, and iii) coalescence rates vary in time since they depend on the structure of the population and the traits present at a given time. Trees can in particular be imbalanced. However, to what extent should these coalescent trees differ from a Kingman’s coalescent is not straightforward since that would certainly depend on the number of lineages. In this section, our goal is to evaluate under which conditions it is possible to detect a statistically significant discrepancy from a Kingman’s coalescent; in other words to what extent does the data allow us to distinguish between the model with interactions and eco-evolutionary feedbacks and a simpler “neutral” model.

We considered statistics commonly used to test the neutrality of the phylogenies of *n* sampled individuals Fu and Li (1993): the number of cherries *C*_*n*_, i.e. the number of internal nodes of the tree having two leaves as descendants, the length of external branches *L*_*n*_, i.e. edges of the phylogenetic tree admitting one of the *n* leaves as extremity, and the time 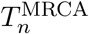 to the most recent common ancestor (MRCA).

We compare the distributions of the normalized *C*_*n*_ and *L*_*n*_ and the distribution of 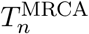 for the forward-backward Doebeli-Dieckmann’s coalescent and the Kingman’s coalescent. Fig. 3.3 shows that these distributions largely differ. For Kingman’s coalescent, asymptotic normality has been established for *C*_*n*_ and *L*_*n*_ (see Blum and François (2005); Janson and Kersting (2011)). The distribution of 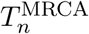 for the Kingman coalescent is computed by using the fact that the trees are binary with exponential durations between each coalescence. We see on Fig. 3.3 that the distribution of external branch length under our model shows an asymmetrical leptokurtic distribution: the external branch length tends to be much shorter than under a Kingman’s coalescent. However, the time to the MRCA is much shorter in Kingman’s coalescent than in our model. The distribution of the number of cherries follows a symmetrical bell-shaped distribution flattened around the mode.

**Figure 3.3:**
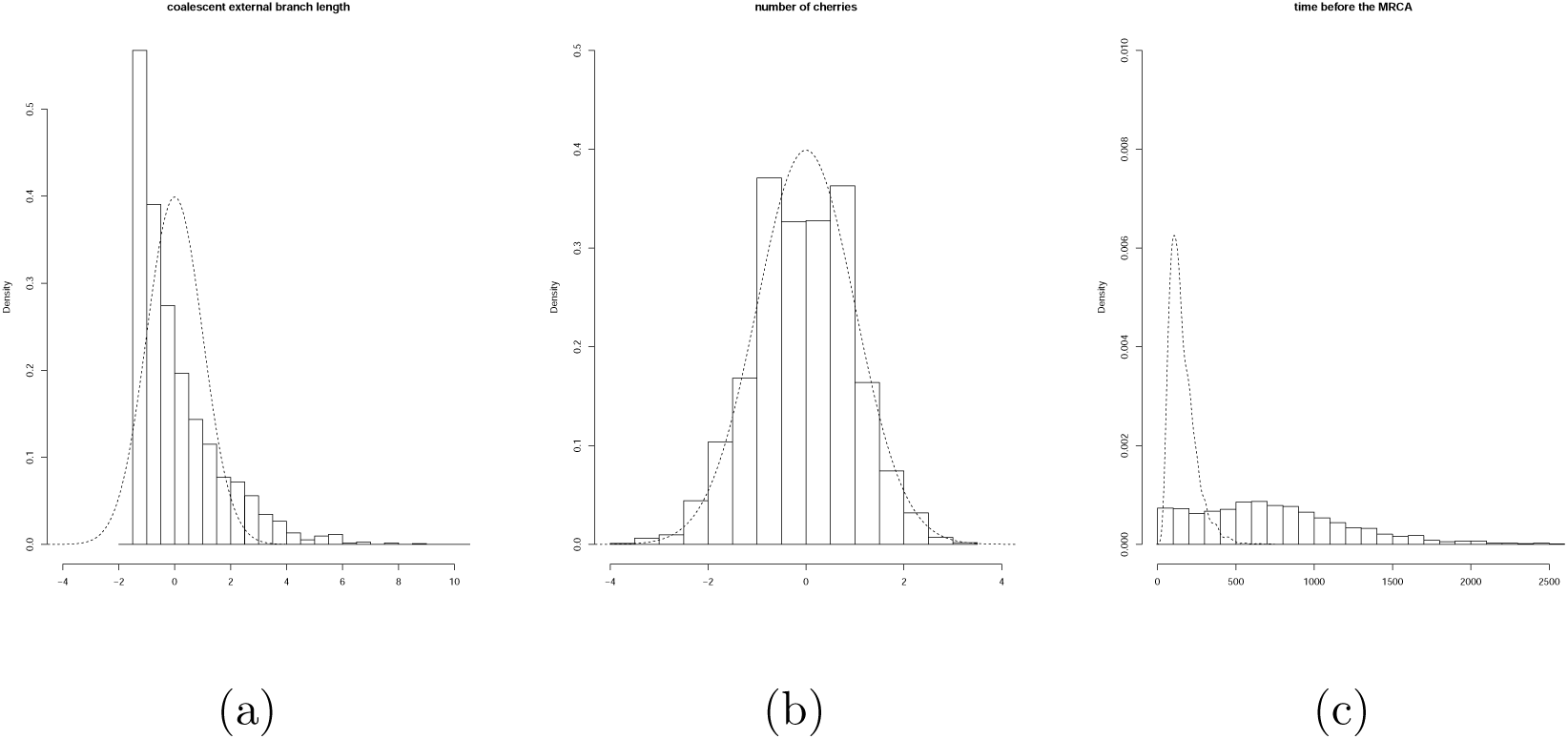
*Histograms of (a) the renormalized external branch lengths, (b) the renormalized number of cherries, (c) the time to the MRCA. The simulations are shown for p* = 0.0076, *q* = 0.7503, *σ*_*b*_ = 1.186, *σ*_*c*_ = 0.4951, *σ*_*m*_ = 0.1448, *η*_*c*_ = 0.0211 *and t*_*sim*_ = 1025.619 *(set of parameter A in Table 1. Results for three other ‘reference’ sets are given in App. E). The dashed line represents the distribution followed by a Kingman’s coalescent (Gaussian distribution for (a) and (b), simulations for (c)).*

We performed statistical tests based on the above mentioned asymptotic normality results for the normalized *C*_*n*_ and *L*_*n*_. We see on the simulations that the number of lineages *m* at the time of sampling plays a role in the shape of the genealogical trees. Thus, we chose to perform neutrality tests conditionally on this number *m*. For each *m*, we chose as pseudo-data one of the simulations of our model with *m* species at the final time, and we performed normality tests for *C*_*n*_ and *L*_*n*_, and an adequation test for the expected distribution under Kingman for 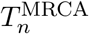. This was repeated 100 times for each value of *m ∈ {*1, … 10*}* (details given in App. E).

Fig. 3.4(a) shows the distributions of the *a posteriori* p-values for the normality tests for *L*_*n*_ and *C*_*n*_. The test for *L*_*n*_ shows that coalescent trees significantly differ from Kingman’s coalescent trees because external branches are shorter in presence of competition between and within lineages as shown in Fig. 3.3(a). Hence, using a Kingman’s coalescent model and ignoring the trait structure of a population tend to overestimate the recent coalescent times. When considering *C*_*n*_, the topology of the neighborhood of the leaves is not always found to significantly differ between Kingman and Doebeli-Dieckmann’s cases, see Fig. 3.4(b) where p-values have a median close to 0.05. There is no systematic acceptance of the Gaussian distribution, even in the case of one single final lineage where multiple merges and varying coalescence rates may create differences with Kingman’s coalescent. Finally, Fig. 3.4(c) shows the distribution of the time to the MRCA depending on the number of lineages. A mean comparison test shows that the mean of the 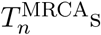 obtained from the simulations of our forward-backward coalescent significantly differs from the expected MRCA time under a Kingman’s coalescent (see App. (E.3)).

**Figure 3.4:**
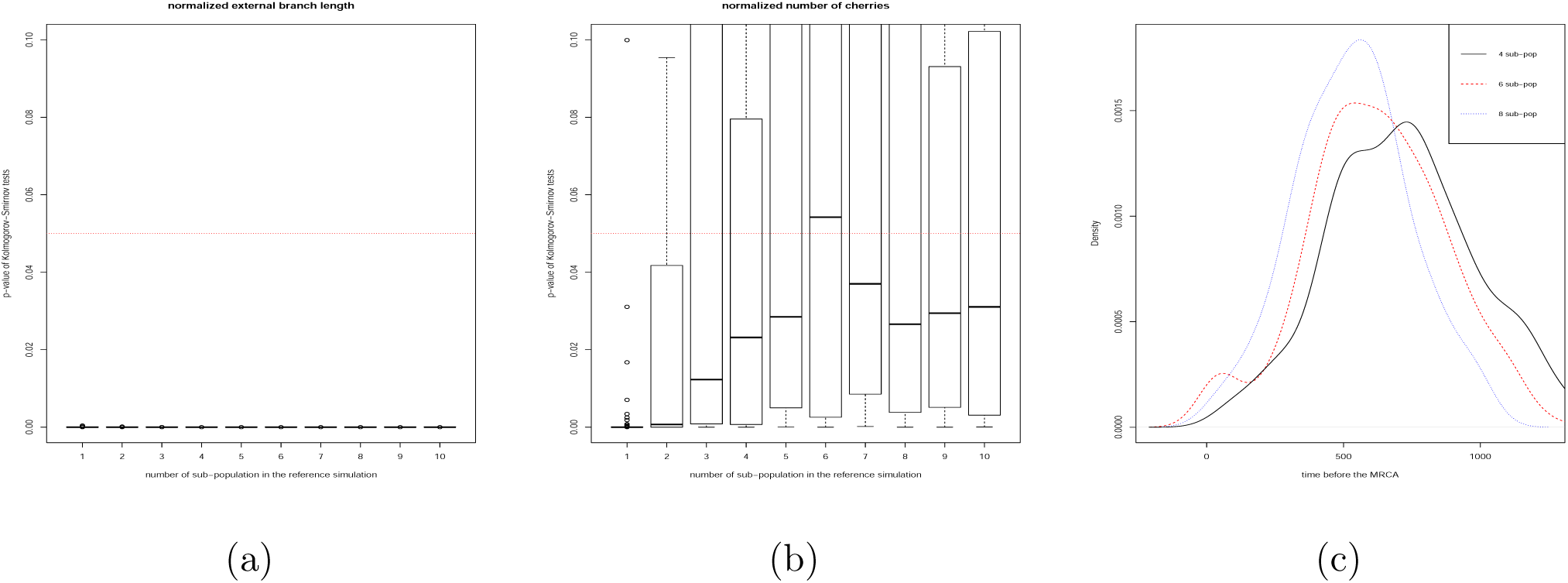
*(a): External branch length L*_*n*_: *Box-plot of the p-values of the Kolmogorov-Smirnov test, for each value of the number of lineages m at sampling time (in abscissa). (b): Number of cherries C*_*n*_: *Box-plot of the p-values of the Kolmogorov-Smirnov test, as a function of m. For (a) and (b)*, 100 *ABC analysis were done for each value of m and we tested if the distribution of the normalized external branch length follows a Gaussian distribution (H*_0_*). The threshold value of rejection of H*_0_, 0.05, *is represented by the dashed red line. If the p-values are lower than this threshold, the distribution of the statistics (L*_*n*_ *or C*_*n*_*) of the forward-backward coalescent trees generated by a Doebeli-Dieckmann model is significantly different than the one under a Kingman’s coalescent. (c): Compared distributions of the age of the MRCA for the forward-backward coalescent (plain line) and for the Kingman’s coalescent (dotted line).*

Overall, we can conclude that the coalescent trees generated by a Doebeli-Dieckmann model significantly differ from a Kingman’s coalescent, as expected, because eco-evolutionary feedbacks and competitive interactions between lineages affect coalescent rates.

### 3.3 Application 2: correlations between genetic and social structures in Central Asia

In Anthropology, a common question is whether or not socio-cultural changes can affect demographic parameters, such as fertility rates. For instance, it is hypothesized that agriculturalists have a higher fertility than foragers (*e.g.* Sellen and Mace (1997)), which is supported by several studies (*e.g.* Bentley *et al.* (1993); Ross *et al.* (2016)). In this section, we analyze genetic data in order to test whether populations with two different lifestyles and social organizations show different fertility rates. Nineteen human populations from Central Asia have been sampled in previous studies (Fig. 3.5(a), Chaix *et al*. (2007); Heyer *et al*. (2015)). Two types of socio-cultural organizations are encountered: Indo-iranian populations are patrilineal, i.e. mostly pastoral and organized into descent groups (tribes, clans…); Turkic populations are cognatic, i.e. mostly sedentary farmers organized in nuclear families. 631 individuals have been sampled (310 from a cognatic population, 321 from a patrilineal one). Ten microsatellite loci have been genotyped on the Y-chromosome. Since there is no recombination on the sexual chromosomes in humans, it is appropriate to use our model which assumes clonal reproduction. Hence, we will perform ABC analysis on the genetic diversity following the paternal lineages.

**Figure 3.5:**
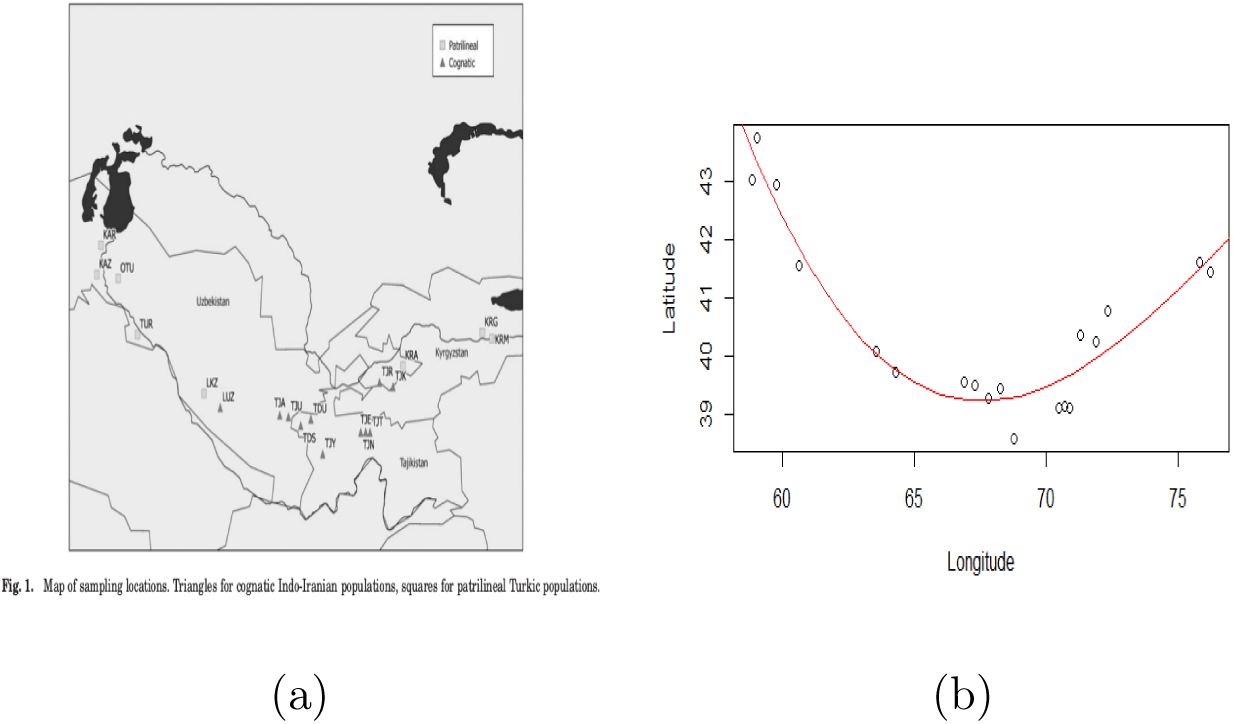
*(a): Map of sampling locations from* *Heyer* et al. *(2015)*. *Triangles correspond to cognatic IndoI-ranian populations, quares to patrilineal Turkic populations. (b): Regression of the data to a 1-dimensional problem.*

We considered that the trait *x* in the model is a vector containing the geographic location of the population and the social organization (cognatic or patrilineal). For geographical positions, given the Fig. 3.5(a), we consider that geographic location is 1-dimensional: we can fit a polynomial curve through the geographical positions of the tribes:

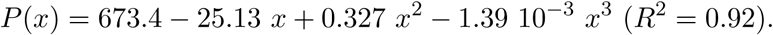

Hence the location of each population is given by the coordinates (*x, P*(*x*)) (Fig. 3.5(b)). The distance between populations is computed thanks to the line integral along the interpolated curve (see details in App. F.2). The neutral marker *u* is a vector containing the genotype at the ten microsatellites.

Our aim is to use our ABC procedure on the genetic data to estimate the parameters *θ* = (*p*_xb01_, *b*_0_, *b*_1_, *p*_loc_, *q, σ*_loc_, *η*_0_, *η*_1_, *σ*_*c*_, *t*_*sim*_) of our model. The individual birth rates is assumed to depend on social organization only and not on geographic location: *b*_0_ for the patrilineal populations and *b*_1_ for the cognatic ones. Death rates are supposed to be due to density-dependent competition for the sake of simplicity: the competitive effect of an individual located at coordinate *y* on an individual in a patrilineal (resp. cognatic) population at location *y*′ is supposed 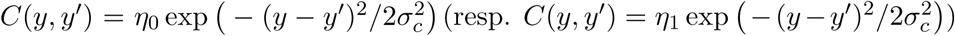. The individual death rate at location *y* is given by the sum of the competitive effects of all individuals. We supposed that individual can found new population after dispersal (corresponding to a mutation on the trait *x* at birth), with probability *p*_loc_, and/or change of social organization, with probability *p*_xb01_. The location of the new population is randomly drawn in a centered Gaussian with standard deviation *σ*_loc_. Following anthropological data, we assumed that social organization changes are unidirectional only from patrilineal pastoral to cognatic farmers populations Chaix *et al.* (2007). *t*_*sim*_ and *q* respectively are the duration of the coalescent and the marker mutation probability.

Estimating the parameter *θ* and using the ABC procedure to select between alternative models will allow us to test whether the null hypothesis

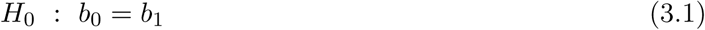

is acceptable, compared to the alternative hypothesis *H*_*a*_ : *b*_0_ < *b*_1_ (see e.g. Grelaud *et al.* (2009); Prangle *et al.* (2013); Stoehr *et al.* (2015)). We generated a set of data with the *a priori* probability 1*/*2 of having *b*_0_ = *b*_1_ and the *a priori* probability 1*/*2 of having *b*_0_ < *b*_1_ (see details in App. F.2). To do this, we generated 10,000 datasets with *b*_0_ = *b*_1_ and 10,000 datasets with *b*_0_ < *b*_1_. The ABC estimation provides weights *W*_*i*_ for each of these 20,000 simulations (see Eq. B.1) that allow to compute the posterior probabilities of each hypothesis: {*b*_0_ = *b*_1_} or {*b*_0_ < *b*_1_}. When the estimated posterior probability for {*b*_0_ = *b*_1_} is smaller than a certain threshold, the null hypothesis *H*_0_ is rejected.

We first checked the quality of the ABC estimation and of the test (3.1) on simulated data (10,000 datasets with *b*_0_ = *b*_1_, 10,000 datasets with *b*_0_ < *b*_1_ where parameters were drawn in their prior distribution). We chose 200 simulations to play in turn the role of the true dataset, 100 among those with *b*_0_ = *b*_1_ and 100 among those with *b*_0_ < *b*_1_. We obtained that parameters estimates were generally close to the true values (App. F.2). After having determined a statistical threshold by minimizing both Type I and II errors, we obtained that the statistical test 3.1 based on the ABC procedure lacked power. This was due to the fact that very often in the simulations, one of the social organization went extinct because of ecological asymmetries: social organization change is supposed unidirectional and *b*_0_ ≤ *b*_1_.

We then performed the statistical test (3.1) on the dataset from Central Asia populations. Using the same ABC framework with the same 20,000 simulations, we computed the posterior distribution conditional on the summary statistics. We then performed the ABC test presented as with the pseudo-data sets (App. F.2). Fig, 3.6 shows the estimated parameters. Our results show in particular that the probability of founding new populations are low (less than 1%) while the probability to change one’s social organization is large (more than 10%). The test (3.1) was found not significant (App. F.3, Fig. 3.7). Hence our result suggest that fertility is not lower in patrilineal than in cognatic populations.

**Figure 3.6:**
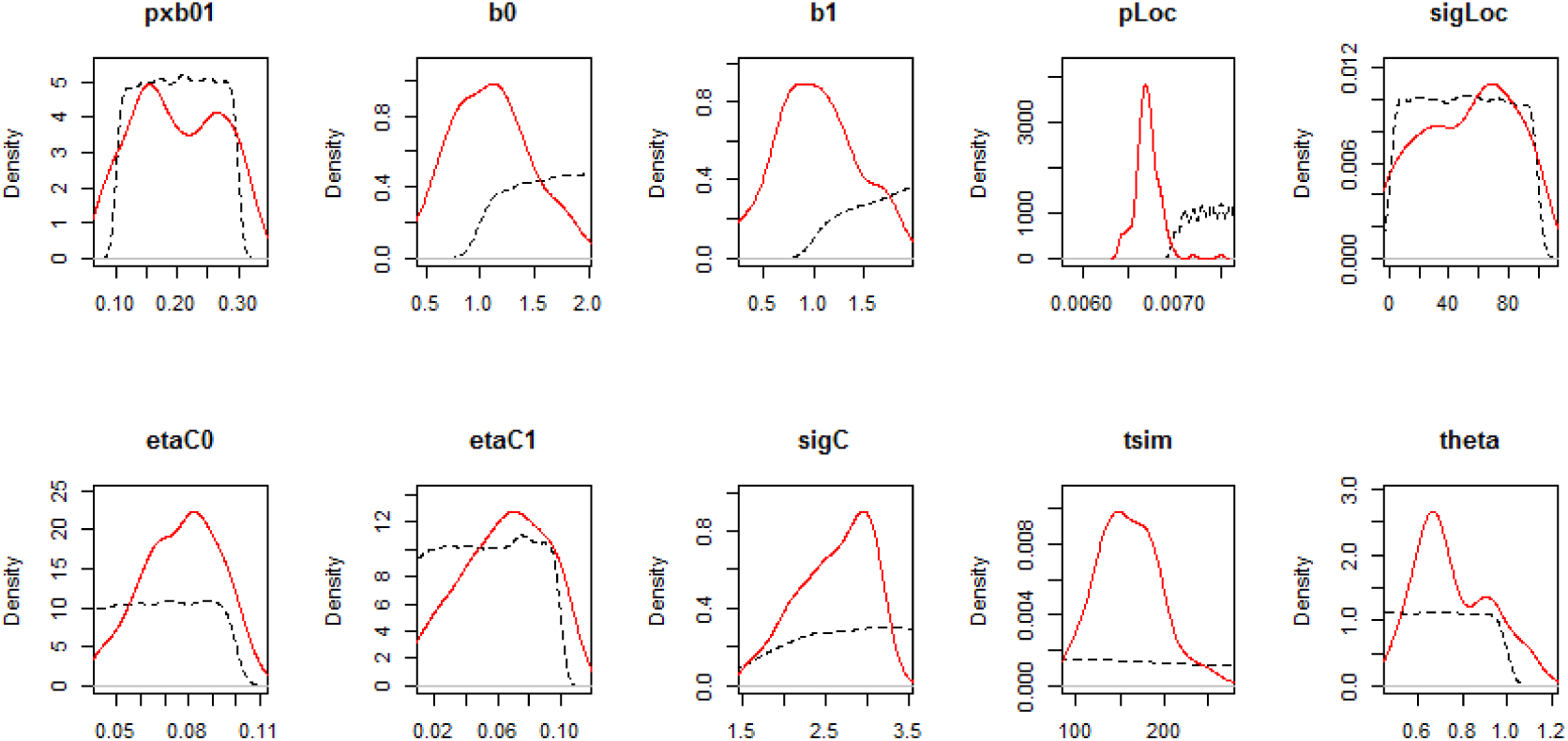
*Results of the ABC estimation for the dataset of Heyer et al.* *Heyer* et al. *(2015)* *for Central Asian populations.*

**Figure 3.7:**
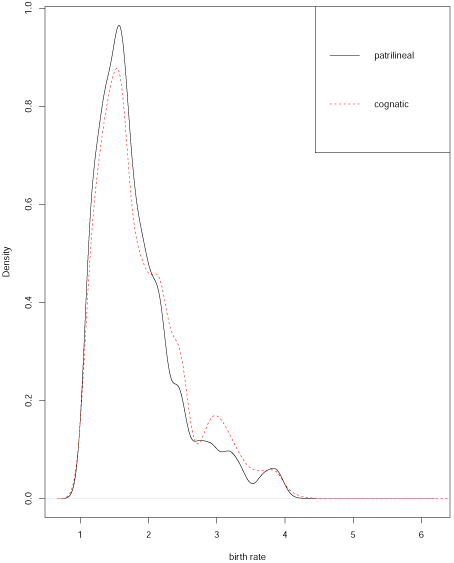
*Approximate posterior distributions for b*_0_ *and b*_1_, *obtained by ABC on the Central Asian database with 20,000 simulations.*

## 4 Discussion

Inferences from genetic data are most often performed under three important assumptions in the existing literature. First, the population size and structure are known parameters: either it is fixed or it follows a deterministic evolution, according to a given scenario (*e.g.* expansion or bottleneck, or a fixed structure with known migration rates between sub-populations). Second, mutations are supposed to not affect the genealogical trees, *i.e.* models are supposed neutral. Selection is rarely explicitly taken into account in inference methods. Third, there is no feedback between the evolution of the population and its demography: a selected mutation is supposed not to affect population size, or the population structure. The most often models used in inference, the Kingman’s coalescent and the Wright-Fisher model, make the three assumptions altogether. The goal of the present paper was to present a model and an inference method which allow to relax all these assumptions. We showed that by using an ABC procedure, it was possible to estimate ecological, demographic and genetic parameters from genotypic and phenotypic data.

Recently, Rasmussen and Stadler (2019) proposed a birth-death model without interactions where mutations can affect the birth and death rates of individuals in a strain, which in return affect the genealogies. They showed how it was possible to use phylogenies to estimate the effect of mutations on fitness in some viruses. In our paper, we go a step further by allowing interactions between individuals, and population structure and demography that depend on the evolution of the population. Our model assumes two genetics traits, a selected trait which governs the structure of the population, and a marker linked to the trait which is neutral and used to infer the genealogy. We first showed how genetic diversity at the neutral marker is related to the evolution at the selected trait, and to the size and structure of the population. We then used this relationship by developing ABC procedure which allows to estimate ecological parameters based on genetic diversity at the neutral marker and on the partial or total knowledge of the population structure. We showed on simulated data that the ABC procedure gives accurate estimates of ecological parameters such as the birth, death and interactions rates, and genetic parameters such as the mutation rate. Our results also showed that non-neutral genealogies can easily be detected under our framework.

We applied our model and its ABC procedure to reanalyze the genetic diversity of microsatellites on Y chromosomes in Central Asian populations. The genetic diversity are compared between two social organizations and lifestyles: patrilineal vs. cognatic. Previous studies showed a significantly different genetic diversity and coalescent trees topologies, which was interpreted as evidence of the effect of socio-cultural traits on biological reproduction, due to how wealth is transmitted within families Chaix *et al.* (2007); Heyer *et al.* (2015). However, these conclusions were obtained under simplifying assumptions: genealogies followed a modified Wright-Fisher model, and the genetic diversity and coalescent trees topologies were compared independently, *i.e.* there was no interaction between populations and between social organization. Such assumptions dismissed the possibility that socio-cultural traits and social organization could change, that new populations can be founded, and that competitive interactions between individuals within and between social organizations might affect demography and evolution. We relaxed all these limitations by applying our model. We supposed that the trait under selection can affect the birth rate. Contrarily to Heyer *et al.* (2015), we did not test whether wealth transmission could explain differences in genetic diversity and coalescent trees topologies. Rather, we addressed a long-standing question in anthropology: can fertility be affected by a change in a social organization, in particular with a change in the agricultural mode. We found no evidence of a fertility difference between both kinds of social organization. Our findings then ask the question why human populations can adopt new socio-cultural traits without any strong evidence of a biological advantage. Further analyses and data would be necessary to confirm our results, especially regarding the number of children per females. In the data, this information is based on a few interviews that are not at all precise (see (Chaix *et al.*, 2007, Table S3 in the suppl. mat.)). However, since the genetic diversity sampled in contemporaneous population is due to long historical process, it seems difficult to estimate fertility for several dozens or hundreds generations. Our results only suggest that there is, on average, no evidence of an effect of a social trait on fertility all along the history of Central Asia populations. Finally, our paper illustrates that it is actually possible to merge ecological and genetic data and models. Our model is based on classical competitive Lotka-Volterra equations, under the assumptions of rare mutations relatively to ecological processes. The genealogies and genetic diversity produced under such a model are then used to infer ecological and demographic parameters. We showed that relaxing strong assumptions of genetic models is possible, and that it allows to provide new analysis methods. The development of stochastic birth and death models, with (this paper) or without (Rasmussen and Stadler (2019)) interactions open the way to new methods for analyzing data. As highlighted by Frost *et al.* (2015), this is particularly important for the study of epidemics and pathogens evolution. These authors give a list of current challenges which can partly addressed thanks to the method and models developed here. For instance, the role of the host structure on the pathogens evolution and genetic diversity, the role of stochasticity, and providing more complex and realistic evolutionary models.

## Acknowledgements

The authors thank Frédéric Austerlitz and Raphaëlle Chaix for discussion and for sharing anthropological data from Central Asia. This research has been supported by the Chair “Modélisation Mathématique et Biodiversité” of Veolia Environnement-Ecole Polytechnique-Museum National d’Histoire Naturelle-Fondation X. V.C.T. also acknowledges support from Labex CEMPI (ANR-11-LABX-0007-01).

## Data archiving

The genetic data, simulation and the programs developed in the paper wll be archived on Dryad and Github.

## A Mathematical construction of the PES and of the forward-backward phylogenies

Recall the stochastic individual-based model described in Section 2.1, which is parameterized by *K* that we will let go to infinity. Because the population has variable size, it is conveniently represented at time *t* by the following point measure:

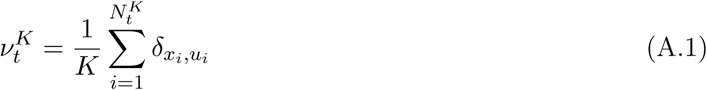

where 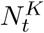 is the size of the population at time *t* and where each individual is represented by a Dirac mass weighting its trait and marker values (individuals being ranked in the lexicographical order, for instance).

In this paper, we denote by ℳ_*F*_(*E*) the space of finite (non-negative) measures on *E*, and by ℳ_1_(*E*) the set of probability measures on *E*. For a non-negative measurable or integrable function *f* on *E* and measure *µ* ∈ ℳ_*F*_(*E*), we define ⟨*µ, f*⟩ = ∫_*E*_ *fdµ*.

In the sequel, we assume that

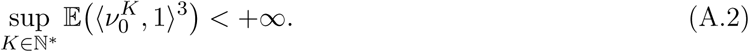

### A.1 Large population limit in the ecological time-scale

First, when *K* → +∞, the mutation rates vanish and we recover in the limit a system of ordinary differential equations.

#### Proposition A.1

*Let us assume that the initial condition* 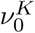 *converges, when K* → +∞, *to a trait-monomorphic initial condition of the form* 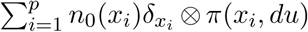, *where p* ∈ ℕ, *x*_1_, … *x*_*p*_ ∈ 𝒳 *and for all i* ∈ {1, … *p*}, *π*(*x*_*i*_, *du*) *is the marker-distribution conditional to the trait x*_*i*_. *Then the marginal trait-distribution of ν*^*K*^ *converges to a limit of the form* 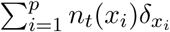 *where* (*n*_*t*_(*x*_1_), … *n*_*t*_(*x*_*p*_)) *are solution of the following system of ordinary equations* (2.2). *The convergence is uniform on every compact time intervals, and holds in probability.*

**Proof** The proof is a standard proof of tightness-uniqueness argument (see Ethier and Kurtz (1986) or Fournier and Méléard (2004)). □

For *p* = 1, we recover the classical logistic equation

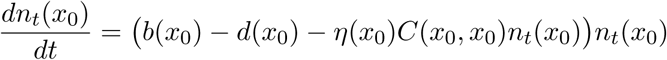

whose solutions started at any non-zero initial condition all converge to the unique stationary stable solution, in case *b*(*x*_0_) − *d*(*x*_0_) > 0:

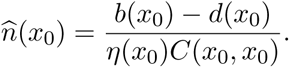

For *p* = 2, we recover a competitive 2 species Lotka-Volterra system, whose solution converges to a stationary stable solution, that can correspond to the extinction of one or both species or coexistence. For polymorphic populations with *p* > 2, the dynamics becomes much more complicated (see Zeeman (1993) for *p* = 3). However the following criterion from Champagnat et al. Champagnat *et al.* (2010) ensures the convergence to a stationary stable point:

#### Proposition A.2 (Champagnat Jabin Raoul)

*Assume that for all i, j* ∈ {1, … *p*}, *η*(*x*_*i*_)*C*(*x*_*i*_, *x*_*j*_) = *η*(*x*_*j*_)*C*(*x*_*j*_, *x*_*i*_) *and that the matrix* (*η*(*x*_*i*_)*C*(*x*_*i*_, *x*_*j*_))_1≤*i,j*≤*p*_ *is positive definite. Then, starting from any initial condition* (*n*(*x*_*i*_; *x*_1_, … *x*_*p*_), *i* ∈ {1, … *p*}) *in the positive quadrant, the Lotka-Volterra system* (2.2) *converges to a unique stationary stable point. In the sequel, we denote this equilibrium* 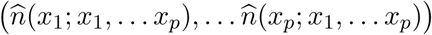.

Notice that the conditions of Prop. A.2 are satisfied for instance when *η*(*x*) ≡ *η* is a constant function and when *C*(*x, y*) = *C*(*x* − *y*) is symmetric with positive Fourier transform. This is in particular the case for Gaussian kernels 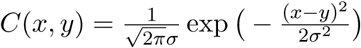. When the competition kernel is symmetric with positive Fourier transform, the equilibria of (2.2) do not depend any more on the initial conditions.

### A.2 Substitution Fleming-Viot limit in the trait-mutation time-scale

Now let us return to the microscopic population (A.1). There are three timescales underlying its dynamics: 1) the ecological timescale of births and deaths is the more rapid, the global birth and death rate being of order *K*; 2) the timescale of marker mutations is the intermediate timescale, when the population size is of order *K*, marker mutations appear at a global rate *r*_*K*_*/K*; 3) the trait mutations happen on the slower timescale; for a population size of order *K*, the global trait mutation rate is of order 1*/K*.

We now consider the dynamics in the timescale of trait mutations and consider 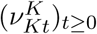. The dynamics of this process, when *K* → +∞, has been studied in Billiard et al. Billiard *et al.* (2015). It is shown that the sequence 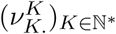 converges to a limit called Substitution Fleming-Viot Process (SFVP) and described as follows. The trait distribution evolves as the Polymorphic Evolution Sequence (PES) introduced by Champagnat and Méléard Champagnat and Méléard (2011). Between trait mutations, the population stabilizes at the equilibrium of the ODE system (2.2) corresponding to the traits which are present in the population. Transitions, whose durations are of order log*K* disappear in the limit. The limiting trait distribution thus jumps from one (possible polymorphic) equilibrium to another one when successful trait mutations arise. Under our assumptions on the mutation probabilities *p*_*K*_ and *q*_*K*_, when a new mutant trait appears and invades into the population, the neutral marker that is linked to it benefits from a hitchhiking phenomenon (see e.g. Barton (1998, 2000); Durrett and Schweinsberg (2004, 2005); Etheridge *et al.* (2006)). Between jumps of the trait distribution, the neutral distribution follows in the limit a diffusive Fleming-Viot process, that boils down to a Wright-Fisher diffusion with mutations in the case where 𝒰 = {*a, A*} has only two elements.

To give a more precise description, let us define the different ingredients appearing in the expression of the SFVP. In a trait polymorphic population at equilibrium, of the form 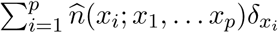, the fitness function of a new small mutant population of trait *y* is defined by (2.3). This fitness function appears in the PES process introduced by Champagnat and Méléard Champagnat and Méléard (2011), defined as follows.

#### Definition A.3

*Let us work under Assumptions of Proposition A.2. The PES process* Λ.(*dy*) *is a pure-jump process with values in* ℳ_*F*_, *the set of point measures on* 𝒳. *It jumps from*

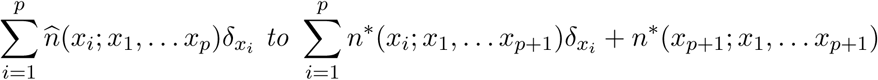

*with rate*

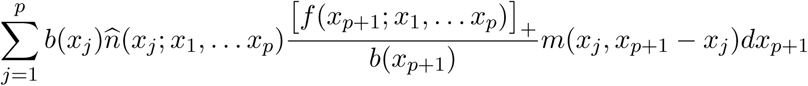

*where the fitness function f has been defined in* (2.3).

By assumptions of Proposition A.2, the sizes *n*\*(*x*_*i*_; *x*_1_, … *x*_*p*+1_) are well defined.

The term 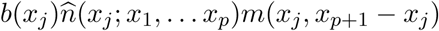 in the jump rate appearing in the definition says that every subpopulation of trait *x*_*j*_ can generate the mutant *x*_*p*+1_, with a probability that depend on the size and birth rate of this subpopulation, and of the mutation kernel. The term [*f* (*x*_*p*+1_; *x*_1_, … *x*_*p*_)]_+_*/b*(*x*_*p*+1_) describes the probability that a mutant trait *x*_*p*+1_ is not wiped out by the stochasticity ruling the births, deaths and competition events.

Finally, let us define the Fleming-Viot process (see Dawson (1993); Dawson and Hochberg (1982); Donnelly and Kurtz (1996) or Etheridge (2000)) that appears in our study.

#### Definition A.4

*Let us fix x* ∈ 𝒳, *u* ∈ 𝒰 *and consider a polymorphic population with traits x*_1_, … *x*_*p*_ ∈ 𝒳 *that is described by the trait-distribution* Λ. *Let us assume that for every continuous bounded test function ϕ on* 𝒰,

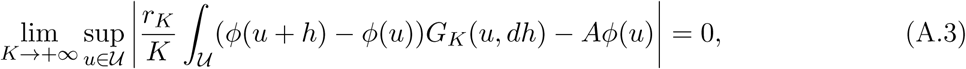

*where* (*A*, 𝒟(*A*)) *is the generator of a Feller semigroup and ϕ* ∈ 𝒟(*A*) ⊆ 𝒞_*b*_(𝒰, ℝ), *the set of continuous bounded real functions on* 𝒰.

*The Fleming-Viot process* 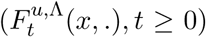 *indexed by x, started at time* 0 *with initial condition δ*_*u*_, *associated with the mutation operator A and evolving in a population whose trait-marginal distribution is* Λ(*dx*), *is the* 𝒫(𝒰)*-valued process whose law is characterized as the unique solution of the following martingale problem. For any ϕ* ∈ 𝒟(*A*),

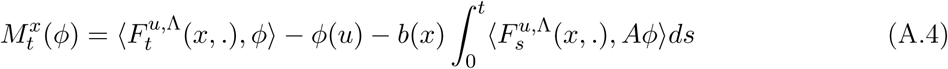

*is a continuous square integrable martingale with quadratic variation process*

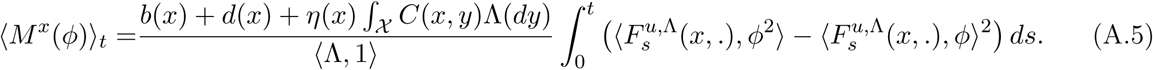

Now we can enonciate the convergence result established in Billiard *et al.* (2015):

#### Theorem A.5 (Billiard, Ferrière, Méléard and Tran)

*Let us assume that* (A.2), (A.3) *and Hypothesis of Proposition A.2 hold.*

*The sequence* 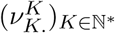 *converges in distribution to the superprocess* 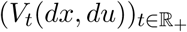 *such that:*

i. *∀t* ∈ ℝ_+_, *V*_*t*_(*dx*, 𝒰) = Λ_*t*_(*dx*), *the PES process of Definition A.3. We write V*_*t*_(*dx, du*) = Λ_*t*_(*dx*)*π*_*t*_(*x, du*), *where π*_*t*_(*x, du*) *is the conditional probability distribution of the marker conditionally to the trait x.*
ii. *At time t, the process V jumps from a state* 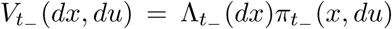 *with* 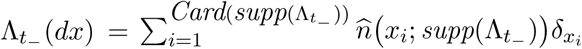 *to*

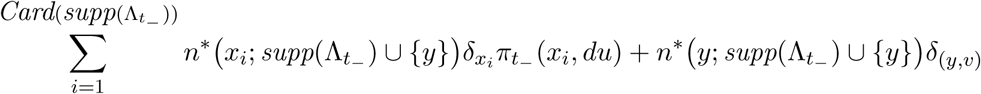

*with rate*

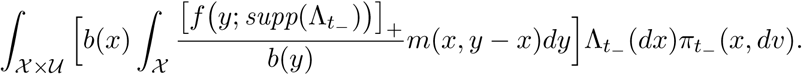 *Let us denote by τ*_0_ = 0 *and τ*_1_, *τ*_2_, … *the successive jump times of V*. *(which are also the jump times of the PES* Λ).
iii. *Between jumps, and conditionally to the trait-distribution, the marker-probability distributions π*_*t*_(*x, du*) *evolve as independent Fleming-Viot superprocesses, characterized by the following martingale problem: for t* ≥ *τ*_*k*_ *(k* ≥ 0*), for x* ∈ *supp*(Λ_*t*_) *and for ϕ*(*u*) ∈ 𝒟(*A*):

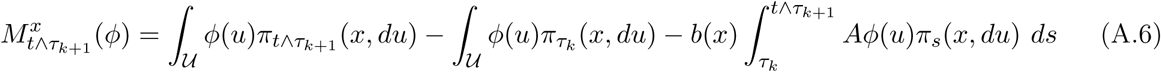

*is a square integrable martingale with quadratic variation process*

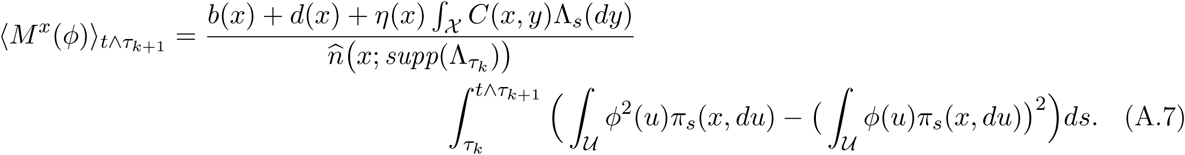 *The convergence holds in the sense of finite dimensional distributions and in the sense of occupation measures.*

Point (ii) tells us that when a new mutant trait appears, there is a hitchhiking phenomenon and the marker that is physically linked with this trait can invade the population (see Billiard *et al.* (2015)). With a similar proof, it can be proved that when there is coexistence of other traits, the marker distributions of the sub-populations corresponding to the traits that coexist after the invasion of the new mutant trait are the same than the marker distributions before the jump.

Heuristically, we can think of the trait *x* ∈ ℝ^*d*^ as characterizing the species to which the individuals belong. Put as this way, the model defined below is a speciation extinction model and we are interested in the phylogenies of a neutral marker in this framework.

### A.3 Phylogeny model for the SFVP: the PES-based phylogenies

In the SFVP limit, the subpopulation of trait *x* living at time *t* say, can be considered of constant “size” 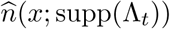, with birth and death rates *b*(*x*) = *d*(*x*)+ ∫_𝒰_ *C*(*x, y*)Λ_*t*_(*dy*). The competition term implies that the death rate in the species *x* - or alternatively the subpopulation size 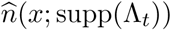 - depends on the composition of the whole population: each time a new species appears or each time a species gets extinct, the sizes of the species populations change.

Between mutations, the marker distribution evolves as a Fleming-Viot superprocess and thus we can expect that the phylogenies of individuals sampled from the species *x* at time *t* are distributed as a Kingman coalescent with rate *b*(*x*) and efficient size 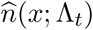 (that depends a priori on the complete set of traits in the population), with a bottleneck at the time *τ*_*x*_ of appearance of the mutant trait *x*.

Let us first recall some results on the phylogenies of Fleming-Viot processes. The links between the processes forward and backward in time requires the notion of duality, and we refer to Etheridge (2000); Jansen and Kurt (2014) for detailed presentations. Two Markov processes 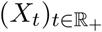 and 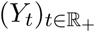 with values in *E* and *F* respectively are in duality with respect to *f* ∈ 𝒞(*E × F*, ℝ), if for every 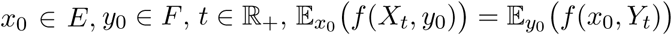. Taking the derivative, this implies that for all *x* ∈ *E* and *y* ∈ *F*, ℒ_*X*_ *f* (., *y*)(*x*) = ℒ_*Y*_ *f* (*x*, .)(*y*), where ℒ_*X*_ and ℒ_*Y*_ are the generators of *X* and *Y*. Applying duality to the Fleming-Viot process, we can obtain the distribution of the phylogenies of a sample on individuals chosen uniformly in the population at time *t*.

#### Lemma A.6

*Let A be a generator and n* ∈ℕ*. *Let ζ*_0_ ∈ *C*(𝒰^*n*^, ℝ) *be a function such that for every i* ∈ {1, …, *n*}, *and for every* (*u*_1_, … *u*_*i*−1_, *u*_*i*+1_, … *u*_*n*_) ∈ 𝒰^*n*−1^, *u*_*i*_ ↦ *ζ*_0_(*u*_1_, … *u*_*n*_) *belongs to* 𝒟(*A*). *Let* 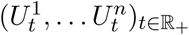 *be a* 𝒰^*n*^*-valued process whose generator G is defined for a function as ζ*_0_ *by:*

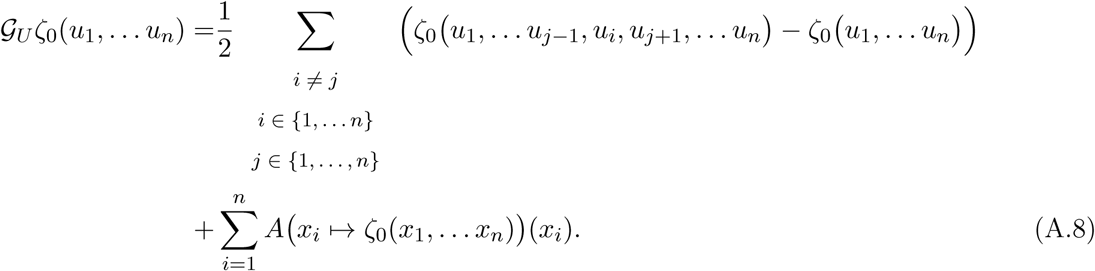

*The genealogies of the process* 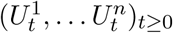 *is a Kingman coalescent with parameter 1.*

**Proof** The generator *G* says that

- for all *i, j* ∈ {1, …, *n*}, the particle *j* is replaced by a particle at the same state as the particle *I* with rate 1*/*2.
- between jumps, each component 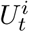 of the vector process evolves independently following the generator *A*.

Then, that the genealogies of the process 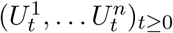 is a Kingman coalescent is straightforward, if we recall that a Kingman coalescent on {1, … *n*} is a process on the set of partitions of {1, … *n*} where two different elements of the partition merge after an independent exponential time of parameter 1. □

#### Proposition A.7

*Let us consider the Fleming-Viot process* (*π*_*t*_(*du*))_*t*≥0_ *started from π*_0_ *and associated with the generator A (we omit here the trait parameters x and the support* Λ *of the trait distribution) that is defined in Th. A.5 (iii). Let n* ∈ℕ* *be the number of individuals drawn independently in π*_*t*_(*du*) *at time t* > 0. *Let ζ*_0_ ∈ 𝒞(𝒰^*n*^, ℝ) *such that for every i* ∈ {1, …, *n*}, *and for every* (*u*_1_, … *u*_*i*−1_, *u*_*i*+1_, … *u*_*n*_) ∈ 𝒰^*n*−1^, *u*_*i*_ ↦ *ζ*_0_(*u*_1_, … *u*_*n*_) *belongs to* 𝒟(*A*). *Let* 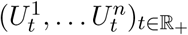 *be a* 𝒰^*n*^*-valued process with generator G defined in* (A.8). *Then*,

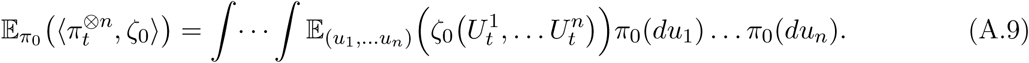

The relation A.9 shows that the distribution of *n* individuals sampled independently from *π*_*t*_ at time *t* is the same as the distribution of the diffusion 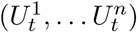 started at independent positions drawn in *π*_0_.

**Proof** First, let us consider the distribution of *n* individuals sampled uniformly in *π*_*t*_(*x, du*), for *x* ∈ 𝒳. To use the duality techniques, we compute 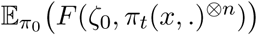 for *ζ*_0_ ∈ 𝒞(𝒰^*n*^, ℝ_+_) and for the test function *F* ∈ 𝒞(𝒞(𝒰^*n*^) *×*ℳ_1_(𝒰^*n*^), ℝ) such that

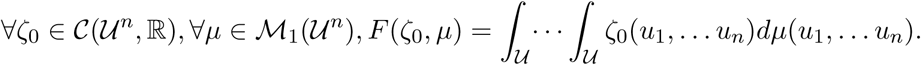

For 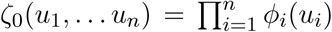 where *ϕ*_1_, … *ϕ*_*n*_ ∈ 𝒞(𝒰,ℝ) bounded, we have for any measure 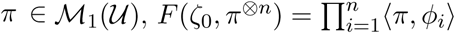. Then, by Itô’s formula for such a function *ζ*_0_,

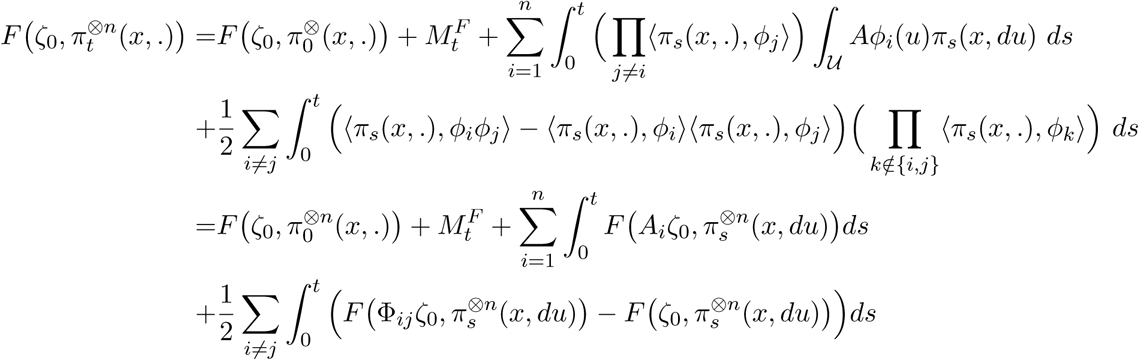

where 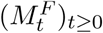 is a square integrable martingale, where *A*_*i*_*ζ*_0_(*u*_1_, … *u*_*n*_) = *A*(*u*_*i*_ ↦ *ζ*_0_(*u*_1_, … *u*_*n*_))(*u*_*i*_), and where Φ_*ij*_*ζ*_0_(*u*_1_, … *u*_*n*_) = *ζ*_0_(*u*_1_, …, *u*_*j*−1_, *u*_*i*_, *u*_*j*+1_, … *u*_*n*_). Taking the expectation:

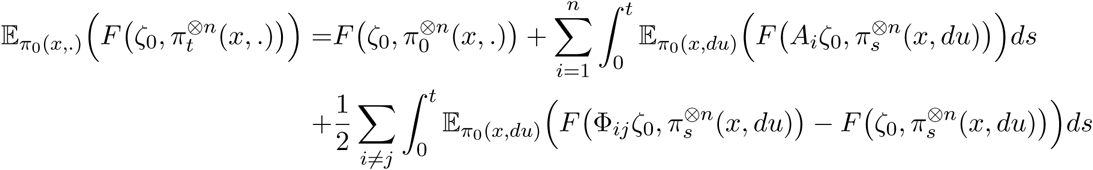

Looking at the generator appearing in the above equation, we introduce the Markov process 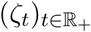 with values in *𝒞* (𝒰^*n*^, R) that jumps from *ζ* to Φ_*ij*_*ζ* with rate 1*/*2 for all *i, j* ∈ {1, … *n*} and whose evolution between jumps is given by the generator 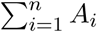. The generator of 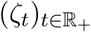 thus satisfies, for a measure *π* ∈ *ℳ*_1_(𝒰^*n*^),

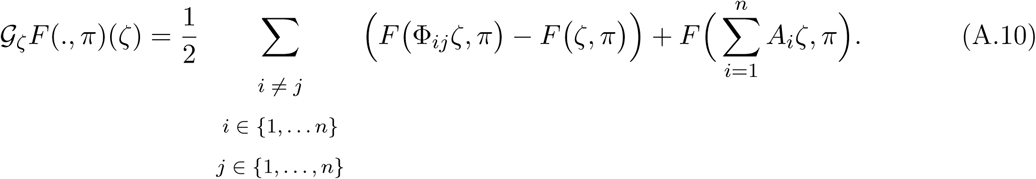

Thus, 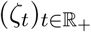 and 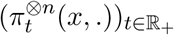 are in duality through *F*, and:

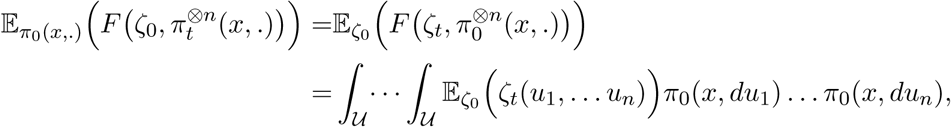

by Fubini’s theorem.

Now, comparing their generators (A.8) and (A.10), we see that 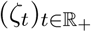 is itself in duality with 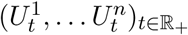 through the function *f* (*ζ*, (*u*_1_, … *u*_*n*_)) = *ζ*(*u*_1_, … *u*_*n*_). Thus:

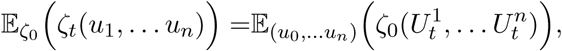

from which we deduce (A.9). □

Thus, it appears that if we draw *n* individuals independently in *π*_*t*_, the distribution of their markers is the same as if we consider a Kingman coalescent started from *n* individuals at time *t*, draw the marker value of the individuals of time 0 independently in *π*_0_ and let these values evolve along the branches according to the generator *G*. In another words, the distribution 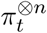 is the same as the distribution of the values of diffusions *G* along the branches of a Kingman coalescent.

From these results, we deduce the forward-backward coalescent: conditionally on the species tree obtained by the PES, we have along each branches a Kingman coalescent whose parameters depend on the sizes of each species, and hence on the entire trait distribution.

### A.4 Simulation of phylogenies with the forward-backward coalescent process

We describe how to simulate the phylogenies of *n* individuals sampled at time 0. Because the simulation starts with obtaining a path of the PES in the forward sense, a parameter −*t*_*sim*_ (*t*_*sim*_ > 0) corresponding to the starting point in the past of the PES is needed. By translation, it is equivalent to say that we start the simulation of the PES at time *t*_0_ = 0 and we reconstruct the lineages of individuals at the present time *t*_*sim*_. Once the PES has been simulated, phylogenies of individuals sampled at time *t*_*sim*_ can be simulated conditionally to the PES, backward time, as described in Section 2.3. We emphasize again that because of competition and interactions between individuals, the phylogenies can not be obtained by considering a sole process that would be backward in time. It would be possible to simulate the phylogenies of the whole population forward time and then to sample within these simulated phylogenies, but that would be time-consuming. Instead, the forward-backward coalescent process is based on the (forward in time) PES, which is a macroscopic process at the population level, and then reconstitutes the phylogenies of sampled individuals at *t*_*sim*_ (backward time).

To keep it intuitive, we describe the algorithm for the Dieckmann-Doebeli model described in Section 3.2.

#### A.4.1 Simulation of the PES

At *t*_0_ = 0, the population is assumed monomorphic. The trait *x*_0_ of the first population is drawn in a normal distribution (mean 0, standard deviation *σ*_*m*_). Its density is the equilibrium of the corresponding isolated population: 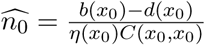.

We then simulate mutation events recursively until the end of the simulation (when time *t*_*sim*_ is reached). Assume that at the current step, the population consists of *p* traits *x*_1_, … *x*_*i*_, … *x*_*p*_. For each sub-population, the duration before the emergence of a possible new mutant in this sub-population is randomly drawn in an exponential distribution of rate 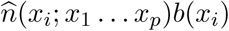.The shortest of these durations determines the mutation that occurs (if its occurs before the end of the simulation). The mutant trait *x*_*mut*_ ∈ [−2, 2] is drawn in a truncated normal distribution with mean the trait value of the sub-population from which it emerges and standard deviation *σ*_*m*_.

Ecological drift can cause the extinction of the mutant before its invasion: its survival probability depends on its initial growth rate in the resident population at equilibrium (the *p* existing populations of trait *x*_1_, … *x*_*p*_ with their densities). If the mutant does not reach a deterministic threshold, which happens with probability [*f* (*x*_*mut*_; *x*_1_, … *x*_*p*_)]_+_*/b*(*x*_*mut*_), nothing happens, and the simulation keeps going with the same traits and densities. Another mutant emergence is then simulated.

Once a mutant emerges, the new Lotka-Volterra system (2.2) must be solved. The mutant density is initially low. Numerically, we took the density 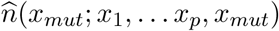 of an isolated population with trait *x*_*mut*_ divided by 100. The system is integrated until it approaches the equilibrium (all growth rates close of 0) and then a Newton method for Lotka-Volterra ODE is used to refine the solution. If a sub-population density is below a certain fixed threshold, it is considered extinct and suppressed from the population.

We store the polymorphic evolutionary sequence (PES): a list of population structures (sub-population traits, densities, emergence time and origin). Example of PES creation and emergence of a mutant is given in Fig. A.1.

#### A.4.2 Simulation of sampled phylogenies

First, *n* = 1000 individuals are sampled in the population at *t*_*sim*_. Their traits depend on the relative abundances of the sub-populations at *t*_*sim*_. If the population consists of *p* species with traits *x*_1_, … *x*_*p*_, the sample is drawn from a multinomial distribution of probabilities

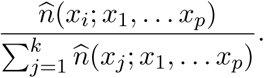

**Figure A.1:**
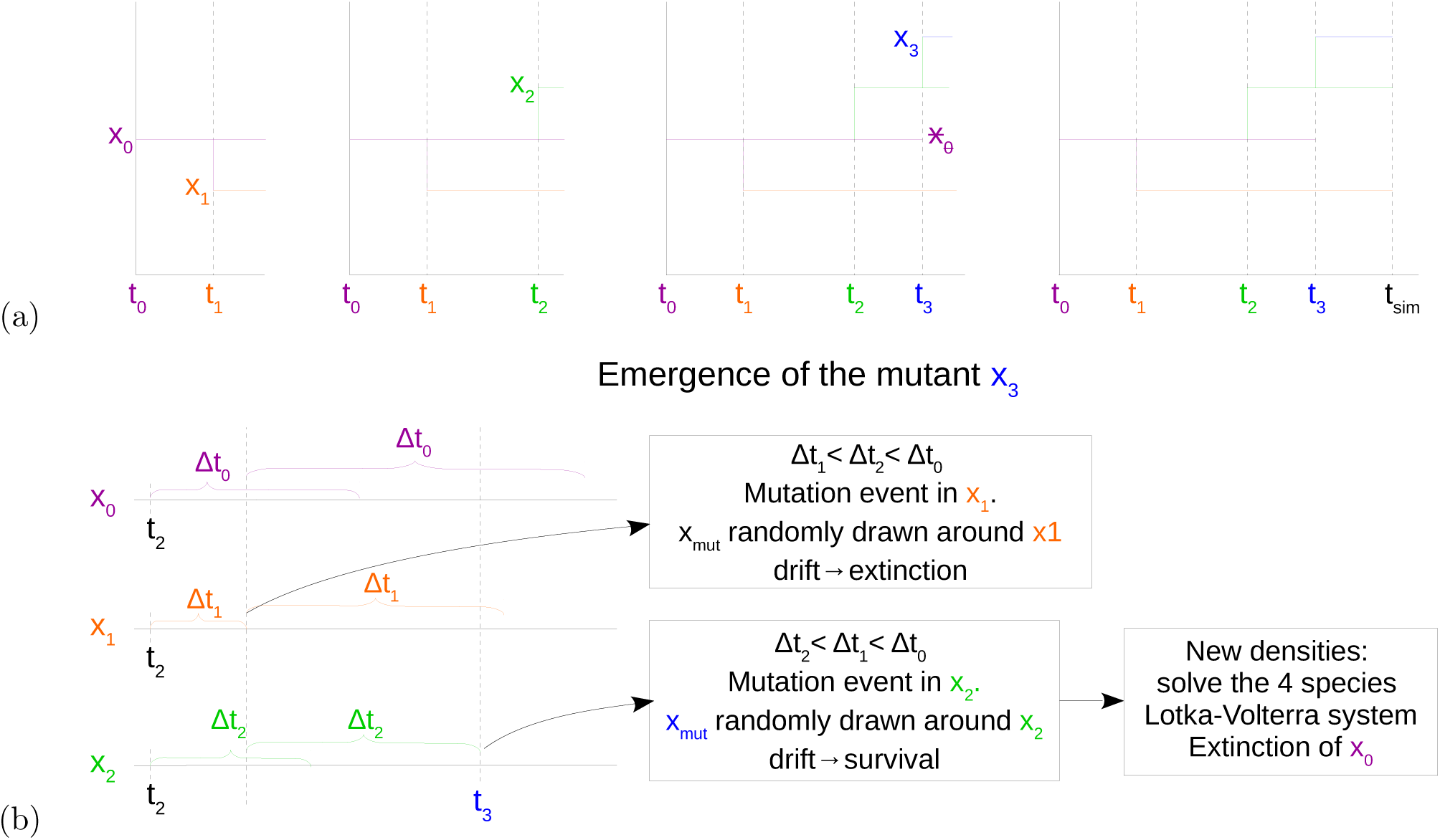
*(a) Example of the creation of a PES. From left: the first mutation time t*_1_ *and the first mutant x*_1_, *which invades, are simulated. Second, a second mutation time t*_2_ *and a second mutant which invade are simulated. Third: a third mutant x*_3_ *invades and the trait x*_0_ *is eliminated by the appearance of this new species. Finally, no more mutation occur before t*_*sim*_. *(b) Example of the emergence of the mutant in a population with three traits x*_0_, *x*_1_, *x*_2_. Δ*t*_*i*_ *corresponds here to the exponential time before the occurrence of a mutant in the species with trait x*_*i*_. *Here the first mutant appears in the population of trait x*_1_ *but fails to invade. Three new clocks are then defined at the time of the failed invasion. It is finally in the species of trait x*_2_ *that the new successful mutant occurs.*

Then, we consider the coalescence of the sampled individuals. Individuals coalesce within their species following the Kingman coalescence model. In a population of *p* species with traits *x*_1_, … *x*_*p*_, to each pair of individual in the same species, say of trait *x*_*i*_ for *i* ∈ {1, … *p*}, is associated an independent random clock with parameter

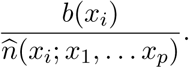

The next coalescence corresponds to the smallest of these exponential random times. The simulation can be done directly at the level of species, which spares some simulation of random exponential variables. In the species of trait *x*_*i*_, the next coalescence hence occurs after an exponential random time of parameter 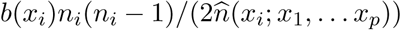; with *n*_*i*_ the number of distinct lineages in the species of trait *x*_*i*_. The two individuals that coalesce are then drawn randomly in the species of trait *x*_*i*_ and are replaced by their common ancestor. Notice that the coalescence rate must be modified at each trait-mutation event.

While going backward in time, we may encounter trait-mutation events. In the sub-population emerging from this mutation event, all individual phylogenies that are still distinct coalesce to produce the common ancestor of the sub-population. This common ancestor is then assigned to its parental sub-population (the sub-population in which the mutant emerged). Then coalescence within sub-population starts again until the next mutation event (or *t*_0_). Coalescence steps are described in Fig A.2.

#### A.4.3 Neutral locus mutation along the phylogeny

If we want to simulate the marker process along lineage, once the phylogenies have been obtained, then we simulate forward in time Brownian motions along each branch of the phylogenetic tree, with every Brownian motion starting from the value at which the Brownian motion of the mother branch stopped.

However, many descriptive statistics of population structure depend on neutral markers rather than on selected traits. This can be handled with a generator slightly different from the one presented in (2.4). We simulated neutral microsatellite markers mutations along the phylogenetic tree: at each mutation event, the marker gain or loss one motif repetition. So here 𝒰 = ℕ and the transitions are *±*1 except if *u* = 0 in which case it can only increase by 1. The mutation events occur in forward time: the mutations are transmitted from the common ancestor to its line of descent until the sampled individuals. When the individual separates into two individuals, they inherit the markers of their parent and then mutations accumulate separately in the two lines.

**Figure A.2:**
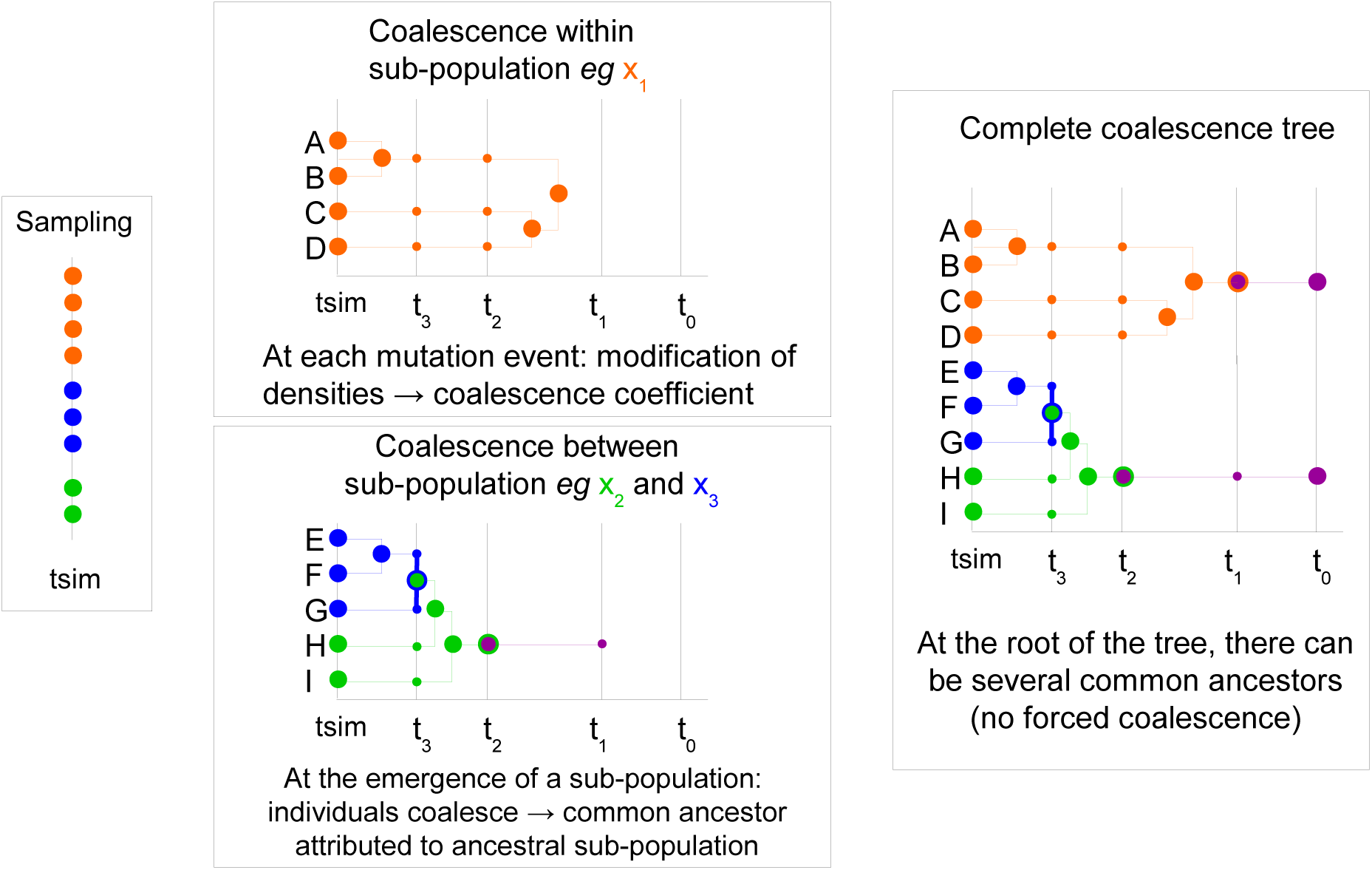
*Example of the emergence of a mutant in a population with 3 traits.*

For each lineage, the time step between each neutral mutation is randomly drawn in an exponential distribution of rate *b*(*x*_*i*_)*θ* (if the trait of the individual changes, the coefficient too). The mutation event results in the loss or in the gain of a motif repeat, with equal probability. Notice that the mechanism generating such marker mutation does not correspond to (2.1). They are much slower than expected. However, the results can be carried to this case: the fact that phylogenies are Kingman coalescent processes can be adapted in this case by adding a color that mutates with the right probability *q*_*K*_.

## B Principle and framework of the ABC estimation

Because the competition and interactions involve the whole population, one is confronted, when trying to write the likelihood of sampled individuals’ phylogenies to formulas where many terms are unobserved. Since we are in continuous time with sub-populations of varying densities, numerical integration over the missing events whose number is unknown is impossible. The ABC method (see Beaumont *et al.* (2002); Marin *et al.* (2012) for a presentation) appears as an alternative which does not use likelihood functions but rather relies on numerical simulations and comparisons between simulated and observed summary statistics. We conducted the ABC analysis using the package abc in R Csillery *et al.* (2012).

The ABC method is a Bayesian method. Let us denote by *θ* the parameters to estimate and by *π*(*dθ*) its prior distribution. The data is denoted by **x**. Instead of trying to estimate the posterior distribution *π*(*dθ* | **x**), which might be very intricate when explicit formulas for the likelihoods are not available, the ABC method proposes to approximate the target distribution *π*(*dθ* | *S*(**x**)) where *S*(**x**) is a vector of descriptive statistics, called summary statistics. Let us denote by *S*_*obs*_ = *S*(**x**) the value of the chosen vector of descriptive statistics computed on the observations.

The principle of the ABC method is to simulate new data corresponding to new parameters drawn into the prior distribution. The parameters that generate data leading to summary statistics close to the observed *S*_*obs*_ are then given a more important weight. The approximation of the target distribution is then the reweighted and smoothed empirical measure of the simulated parameters. The idea of ABC is grounded on non-parametric statistical theory (see e.g. Blum Blum (2010)) and on corrections methods (see Beaumont *et al.* (2002)). More precisely, the ABC estimation requires the following steps:

1. Draw *N* independent parameter sets *θ*_*i*_, *i* ∈ {1, … *N*} in the prior distribution *π*(*dθ*),
2. For each *i* ∈ {1, … *N*}, simulate a realization of the forward-backward coalescent process following Section A.4.
3. Determine for each simulation the corresponding descriptive statistics *S*_*i*_,
4. Compare the descriptive statistics *S*_*i*_, *i* ∈ {1, … *N*}, with *S*_*obs*_ to determine the set of parameter values (*θ*_*i*_) leading to the best fit between *S*_*i*_ and *S*_*obs*_:
  a. rejection method: retain only the *δ* = 0.1% simulations with the lower Euclidean distance between *S*_*i*_ and *S*_*obs*_, by setting a weight *W*_*i*_ = 1*/*(*δN*) to the latter and *W*_*i*_ = 0 otherwise.
  b. smooth re-weighting: to each simulation is computed a weight *W*_*i*_ = *K*_*δ*_(*S*_*i*_ − *S*_*obs*_) where *δ* is a tolerance threshold and *K*_*δ*_ a smoothing kernel.
5. the posterior distribution is then estimated thanks to the weights (*W*_*i*_)_1≤*i*≤*n*_ defined above:

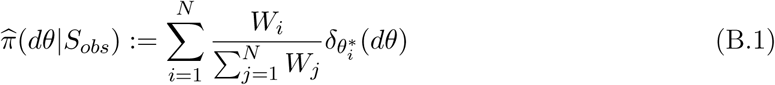

where 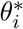 is a correction of *θ*_*i*_ accounting for the difference between *S*_*i*_ and *S*_*obs*_, which we detail in the next paragraph.

In the abc package, several correction methods are available for computing 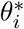. These corrections have been proposed in Beaumont *et al.* (2002). The idea is that if locally around *S*_*obs*_ the parameter *θ* is a function of the summary statistics, say *θ* = *f* (*S, ε*) where *ε* is a noise component, then when *S*_*i*_ is close to *S*_*obs*_, it is possible to correct *θ*_*i*_ = *m*(*S*_*i*_) + *ε*_*i*_ into 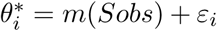 (see also the remark in (Blum and Tran, 2010, Eq. (4.1))). Using the neural network method in the abc package, and based on the couples (*θ*_*i*_, *S*_*i*_), *i* ∈ {1, … *N*}, we can estimate the (possibly non-linear) regression function *m* in the neighborhood of *S*_*obs*_. If *m* is assumed locally linear at *S*_*obs*_, a local linear model can be chosen (loclinear in the abc function), else, neural networks (neuralnet) can be used to account for the non-linearity of *f*. Once the regression is performed, using the estimator 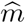 of *m*, the corrected values 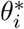 are defined as

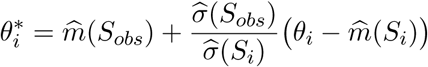

here the bracket in the r.h.s. is an estimator of the residuals *ε*_*i*_. Additionally, a correction for heteroscedasticity is applied (by default) where 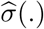 is an estimator of the conditional standard deviation (see Blum and François (2010) for details).

## C Parameters and priors for the estimation of the Dieckmann-Doebeli model

In order to use the ABC method, *N* = 400000 simulations were run using independent random parameter values that are sampled from their prior distributions. The prior distributions of each parameter of the model should reflect the expected values of the parameters. The parameter sets for the four *pseudo-data* sets used in the ABC procedure are given in Tab. 1. Notice that the pseudo-data A and B (resp. C and D) in Tab. 1 have been obtained by simulations with the same sets of parameters. For most parameters, we do not have insights on the expected distribution, thus we choose uniform distributions (details in table C.1).

## D Summary statistics for the ABC estimation based on phylogenetic trees

We detail here several summary statistics that can be used for the ABC estimation. Depending on how much information we have in the data, we can use all these statistics for the ABC or only a subset of them. The three first families (Sections D.1-D.3) provide description of the trait and marker distribution in the population at the final time. These statistics can be computed only at the sample level or also at the population level if additional knowledge is available. Many of these statistics are listed in Pudlo et al. Pudlo *et al.* (2016). The family D.4 contains statistics describing the shape of the phylogenetic tree and can be used if some additional statistics, for instance related to fossils or datation of past speciation events, are available.

**Figure C.1:**
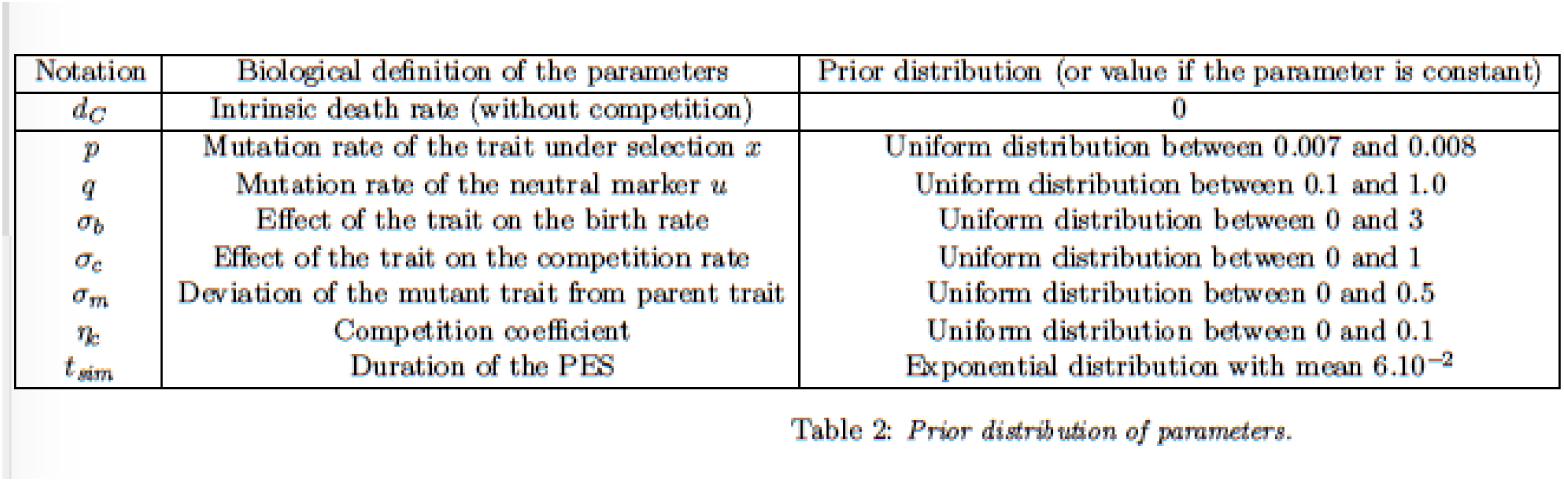
*Prior distribution of parameters.*

### D.1 Population structure from the total population

Statistics measured on the last step of the PES (at *t*_*sim*_):

- number of coexisting species (number of traits in the population),
- mean abundance of the coexisting species,
- variance of the abundance of the coexisting species,
- mean trait *x* of the species,
- trait variance between species,
- mean trait *x* of the individuals (depends on sub-population densities),
- variance between the traits of individuals (depends on sub-population densities).

### D.2 Population structure from the sampled individuals

Statistics measured on the *n* sampled individuals:

- number of sampled sub-populations,
- relative abundance variance between sampled sub-populations,
- mean trait of sampled sub-populations,
- trait variance between sampled sub-populations,
- mean trait of sampled individuals (depends on relative abundance),
- trait variance between sampled individuals (depends on relative abundance).

### D.3 Description of the allelic distribution of the trait and marker

- number of distinct alleles for the marker,
- variance of the marker’s allele distribution,
- genetic diversity (measuring the width of the marker’s allele support). In case the marker corresponds to microsatellites, the genetic diversity is expressed as

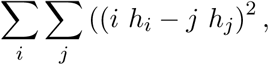

with *h*_*i*_ and *h*_*j*_ the frequencies of the populations with *i* and *j* repetitions. This index corresponds to the expected difference between each couple of individuals in the population.
- Unbiased Gene Diversity, which in the case of microsatellites rewrites as

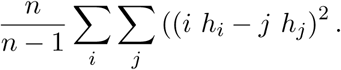 It is the average difference between two individuals in the population.
- M-index: ratio of the number of alleles for the marker on the width of the marker’s allele support,

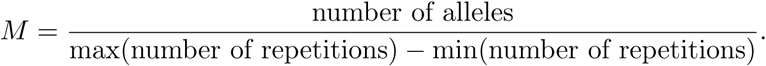 This index is the average percentage of intermediate allelic states that are occupied. The lower the M index, the more the population has lost possible alleles.
- number of traits in the population,
- mean and variance of traits
- abundance distribution for traits.

The allelic diversity is useful to infer population size through time, because drift leads to the loss of alleles in small populations. The M-index is useful to detect reductions in population size (their intensity and duration): during a bottleneck, alleles are lost randomly (not specifically at the end of the range size), thus the percentage of occupied intermediate allelic states decreases (in huge population, alleles should exist throughout the range size). Low allele diversity and low M-index indicate recent reduction of size while low allele diversity but high M-index corresponds to populations that have been small for a long time.

Usually markers that are used correspond to several loci. In this case, additional indices can be added to describe the joint distribution of the marker on these loci. Let us denote by *r* the number of loci. Let us consider two species of traits *x* and *y* respectively. We denote by *x*_*ij*_ and *y*_*ij*_ the frequencies at the locus *j* of the allele *i* in the populations *x* and *y*:

- 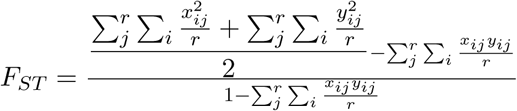
- 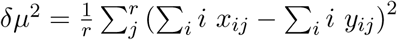. The alleles (*i*) must be expressed as a number of motif repeats.
- Nei’s *D*_*A*_ distance: 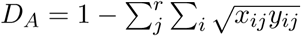
- Nei’s standard genetic distance 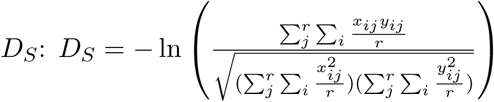

*F*_*ST*_, *δµ*^2^, *D*_*A*_ and *D*_*S*_ are calculated with equal weight given to each population (densities of the populations are not taken into account), or weighted by the densities of the populations (for each pair of population *x* and *y*, the statistic is weighted with 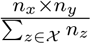.

### D.4 Statistics measured on the PES and on the coalescent tree

Statistics measured on the whole PES:

- number of mutation events along the PES
- Sackin’s index 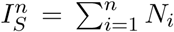 with *n* the number of leaves of the tree and *N*_*i*_ the number of internal nodes from the leave to the root.
- sum of the branch length
- sum of the external branch length

Statistics measured on the coalescent tree:

- sum of the branch length
- sum of the external branch length
- number of cherries (*i.e.* leaves that coalesce together)
- time before the most recent common ancestor
- time before the most recent common ancestor if time of coalescence was neutral

### D.5 Results of the ABC estimation for the sets of pseudo-data

Prior and posterior distribution of parameters estimation determined with the ABC method, for each of the models A-D corresponding to Table 1. All descriptive statistics (blue), statistics with complete knowledge of the population (pink), statistics on sampled individuals, with a knowledge of the trait (red) and statistics on sampled individuals, without any knowledge on the trait (orange).

**Table 1:** *Parameters of the simulations used as data set (referred with their ID in the text)*.

| parameter | ID | $p$ | $q$ | $\sigma_b$ | $\sigma_c$ | $\sigma_m$ | $\eta_c$ | $t_{sim}$ |
| --- | --- | --- | --- | --- | --- | --- | --- | --- |
| value | A | 0.0076 | 0.7503 | 1.186 | 0.4951 | 0.1448 | 0.0211 | 1025.619 |
| value | B | 0.0076 | 0.7503 | 1.186 | 0.4951 | 0.1448 | 0.0211 | 1025.619 |
| value | C | 0.0071 | 0.6815 | 1.7725 | 0.3866 | 0.1372 | 0.0306 | 335.2116 |
| value | D | 0.0071 | 0.6815 | 1.7725 | 0.3866 | 0.1372 | 0.0306 | 335.2116 |

**Figure D.1:**
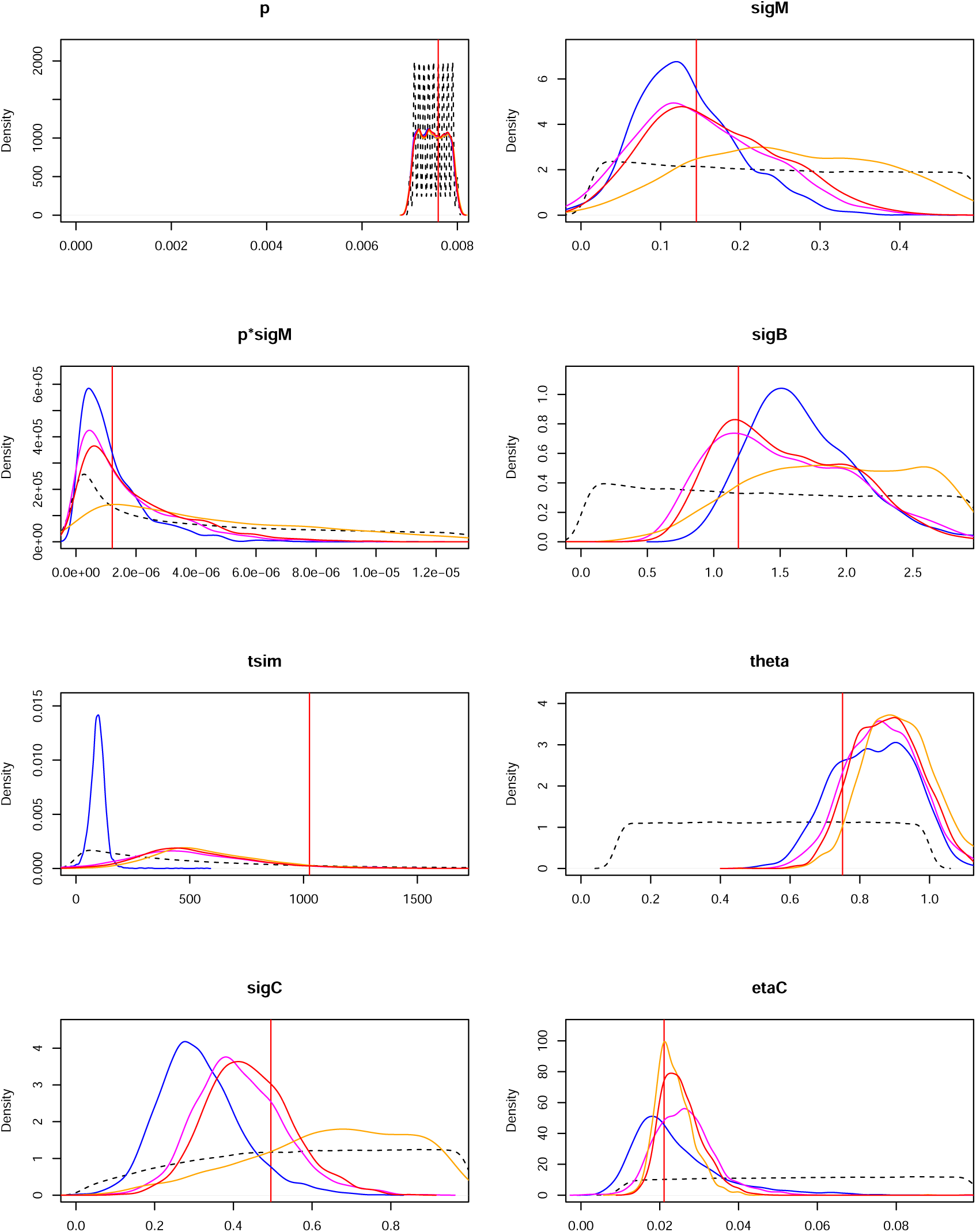
*Results of the ABC estimation for the pseudo-data B (see 1)*

**Figure D.2:**
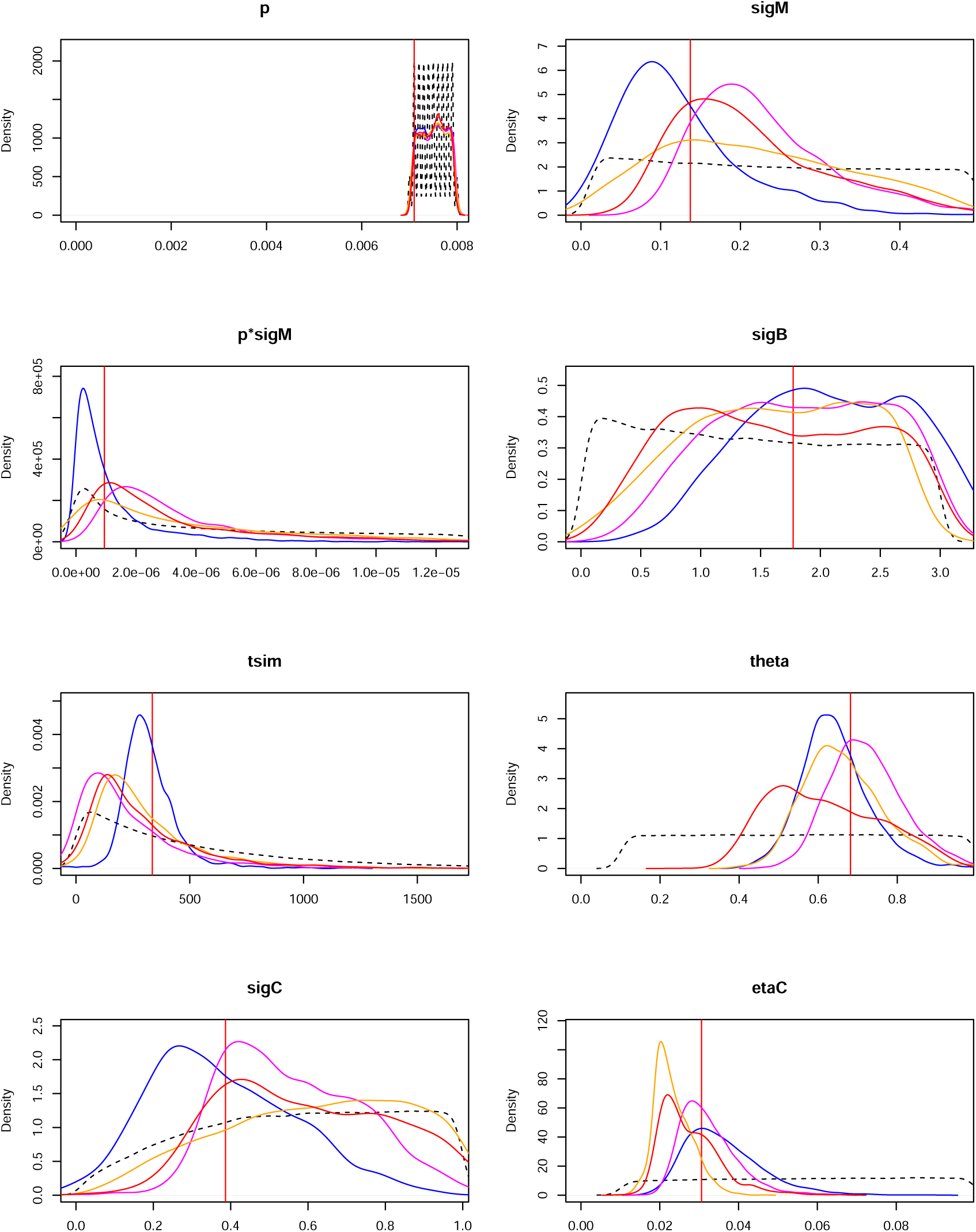
*Results of the ABC estimation for the pseudo-data C (see 1)*

**Figure D.3:**
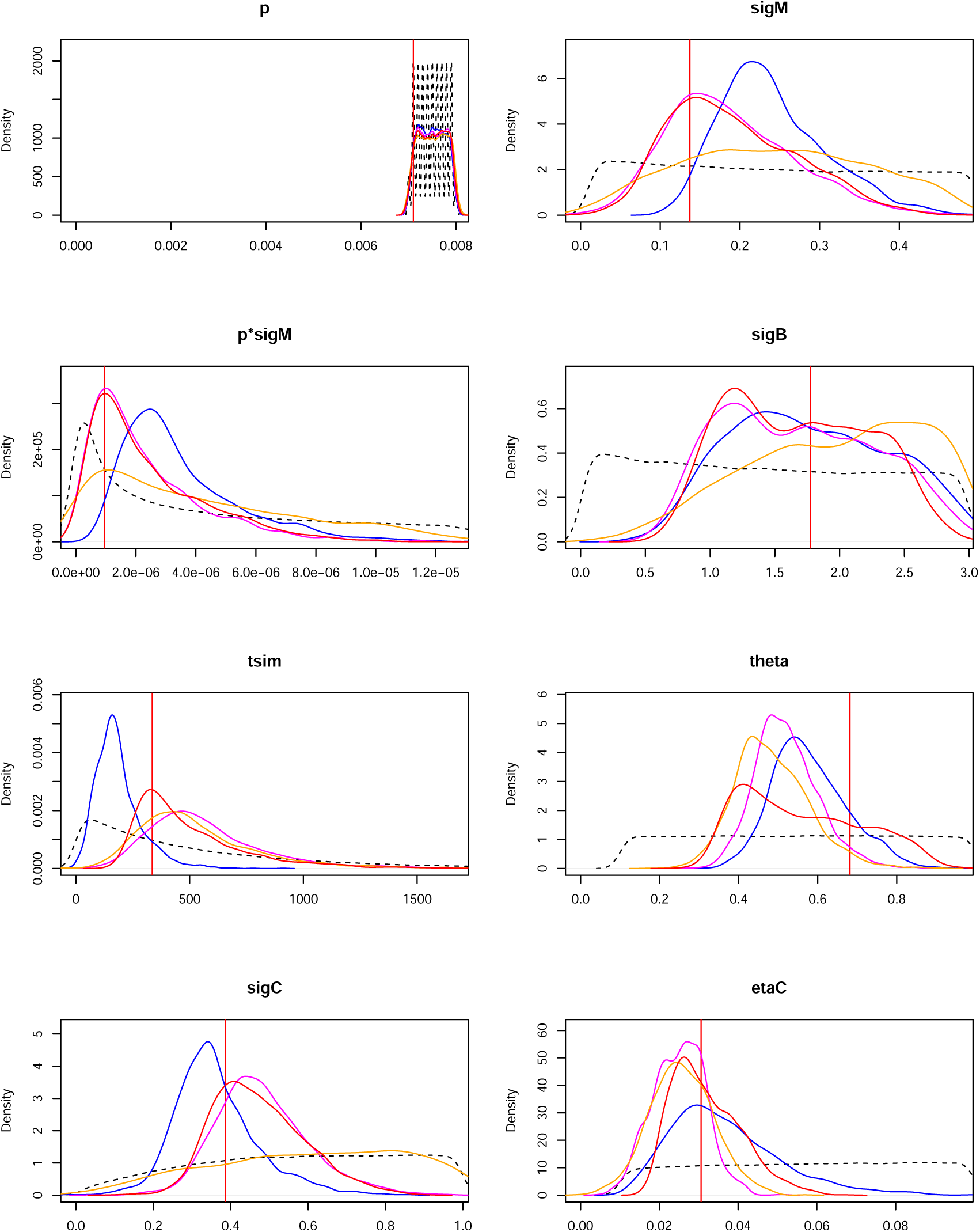
*Results of the ABC estimation for the pseudo-data D (see 1)*

## E Does the Doebeli-Dieckmann coalescent follow a Kingman’s coalescent?

### E.1 Neutrality tests: statistics and procedures

When there is no interaction and when the species play no role, the phylogenies of *n* individuals are described by Kingman coalescent processes. In absence of interaction but when species still have to be taken into account, we have Kingman coalescent processes nested in the species tree. In the latter case, there is no coalescence between phylogenies belonging to distinct species until the speciation event where both species are reunited into a mother species.

In many studies, phylogenies are assumed to stem from neutral models (Kingman coalescent, branching Brownian motions). Then likelihood methods based on this assumption are used for example. The forward-backward coalescent process that is used here brings more complexity but proposes a way to take into account the interactions between species and the fact that the phylogenetic trees might be imbalanced. A first natural step is to check that the new model produces trees that could not be seen as generated by usual models (our choice goes here to Kingman coalescent). This shows that structure has been taken into account.

#### Test statistics

Following the path opened by Fu and Li Fu and Li (1993), we can consider several statistics that are usually chosen as test statistics to test the neutrality of a phylogenetic tree:

- the distribution of cherries, i.e. the number *C*_*n*_ of internal nodes of the tree having two leaves as descendants. The normalized distribution of the number of cherries for a Kingman coalescent follows a Gaussian distribution (see Blum and François (2005)):

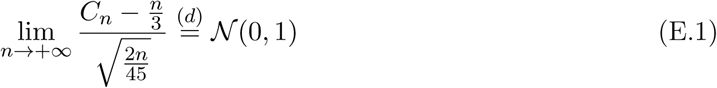
- the length *L*_*n*_ of external branches, i.e. edges of the phylogenetic tree of the *n* sampled individuals admitting one of the *n* leaves as extremity. A beautiful result by Janson and Kersting Janson and Kersting (2011) shows that when *n* converges to infinity, the distribution of the normalized total length of external branches of a Kingman coalescent converges to a Gaussian distribution:

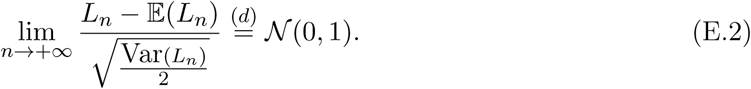
- the time 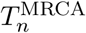 to the most recent common ancestor (MRCA). In a Kingman coalescent, this time is distributed as the sum of *n* independent exponential random variables with respective rates 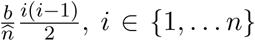, where 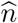 is the density at equilibrium and *b* is the natural growth rate when there is no species structure in the population.

Using the three test statistics that we have presented, we tested whether the observed phylogenies could be described by a Kingman coalescent or not. For the number of cherries and for the external branch length, we computed the renormalized statistics are performed normality tests: a Mann-Whitney test (using the function wilcox.test in R, with unpaired distributions), a Shapiro test and a Kolmogorov-Smirnov test. For the time to the MRCA, we performed an adequation test between the distribution of the MRCA in our model and the distribution of the sum of independent exponential random variables described above.

In all the tests, we set the significance level to *α* = 5% and the null hypothesis *H*_0_ is that the observed tree can stem from a Kingman coalescent. If the p-value of the test is lower than this threshold *α*, then we must reject *H*_0_ and consider that distributions are different in our model and Kingman’s model.

To check that these tests perform well, we applied them to simulated Kingman coalescent processes and checked when the null hypotheses were correctly accepted (see App. E.2).

#### Application to the forward-backward coalescent

To test that the forward-backward coalescent produces non-neutral phylogenies, a first idea would be to perform a Monte-Carlo test by simulating several trees from this model and running the neutrality tests on these simulations. However, such result would be dependent on the parameters chosen for the simulations and would also necessitate a Monte-Carlo loop for exploring the set of parameters. Keeping these ideas in mind, we proceed a bit differently. Assume that we had data **x** generated by the forward-backward coalescent and *a priori* distributions for the parameters *θ*. In this Bayesian framework, would the null hypothesis *H*_0_ : “a phylogeny produced by the forward-backward coalescent is distributed as a Kingman coalescent” be accepted or not?

Given the data **x**, the *a posteriori* distribution of *θ* is computed using ABC, yielding an *a posteriori* distribution on the phylogenies conditionally to **x**. The estimation of the probability ℙ(*H*_0_ is accepted | **x**) can then be approximated by Monte-Carlo using the *N* simulations performed by the ABC procedure: we use the simulations retained by ABC in order to reject abnormal simulations, and the associated weights produced by the ABC. This provides an approximation of the distribution of the test statistics (E.1) and (E.2) conditionally on *S*_*obs*_ = *S*(**x**) yielding in turn an approximation of ℙ (*H*_0_ is accepted | *S*(**x**)):

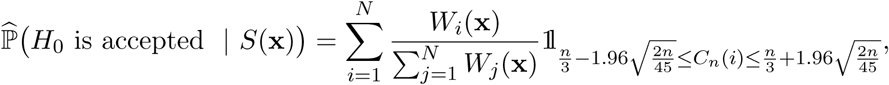

where *C*_*n*_(*i*) counts the number of cherries in the simulation number *i*.

Since this question is investigated in a general framework without data, we use a Monte-Carlo approach to sum over the possible data **x**. Averaging this result over datasets **x** conditionally on the number of species at the final time of the simulations allows to estimate the probability of accepting *H*_0_ conditionally on the number of species at the sampling time. We condition on the number of species at the sampling time as we noticed that this variable impacts the outcome of the test. When there is only one species for example, our model predicts that the phylogenies look like Kingman coalescents (with a possible multiple merge at the first adaptive jump encountered). However, with a growing number of species, we expect a deviation from the Kingman model due to forced coalescence at the creation of each species, and to the different coalescence rates between species.

- for each number of species (*m* from 1 to 10), we randomly chose a simulation that we used as pseudo-data **x** in the ABC an Alyssa. We repeated this action 100 times (providing **x** = **x**_1_, … **x**_100_ for each value of *m*).
- for each **x**, an ABC is performed and provides an approximated *a posteriori* empirical distribution conditionally on *S*(**x**): to each simulations *i* ∈ {1, … *N*} is associated a weight *W*_*i*_(**x**), see (B.1). For each of these weighted simulations, the statistics *C*_*n*_ and *L*_*n*_ can be computed and the normality tests (E.1) and (E.2) can be performed. We deduce from this the approximated *a posteriori* distribution of the p-values.

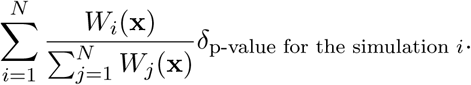
- Averaging over **x**, conditionally on the number of species at the sampling time, we obtain an estimator of the probability that *H*_0_ is rejected conditionally to the number of species at the sampling time.

The normality of the test statistics (E.1) and (E.2) are performed on the same ABC runs. Fig. 3.3 gives the distributions of the normalized external branch length and the number of cherries of the weighted simulations from the ABC analysis presented in the previous section.

We set the significance level to *α* = 5% for all these tests. If the p-value of the test is lower than this threshold, then we must reject *H*_0_ and consider our distribution does not follow a Gaussian distribution. The results for the external branch lengths and cherries are shown in Fig. 3.4(a) and (b) and reveal that our coalescent trees differ from Kingman coalescent trees.

We also wanted to determine whether the coalescence time depended on the number of species. For this, we made a classical test of mean comparison associated with the null hypothesis: *H*_0_ : “the mean time to MRCA in the data is equal to the mean time to MRCA in a Kingman coalescent”. This test is based on the following Student statistic:

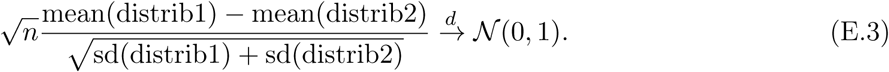

One can see in Fig 3.4(c) that the distributions differ, which is confirmed by the test. This confirms the finding obtained with the study of external branch lengths: timescale suffers from ignoring the interaction between species and can lead to false datings.

### E.2 Neutrality of the model without mutation

We first tested the consistency of our model with Kingman coalescent trees when no mutation of the trait can emerge: the population is monomorphic and the trait mutation is set to 0. We simulated trees with no mutation of the trait and tested the normality of the distribution of normalized external branch length and the normalized number of cherries. We also tested the adequation between the distribution of the time to the MRCA in our model and in a Kingman coalescent. The empirical distributions of these three statistics are represented in Fig E.1. Visually, these empirical distributions fit the targeted distributions under *H*_0_ that are given in Section E.1.

**Figure E.1:**
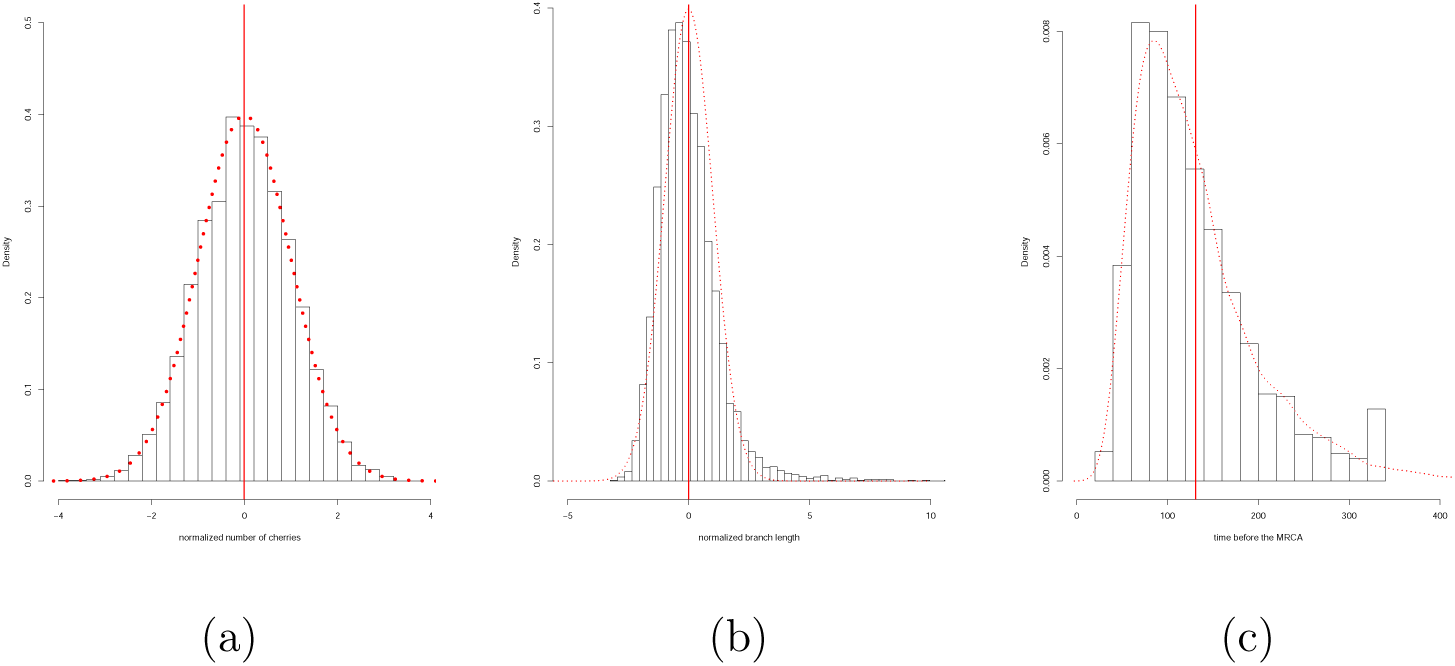
*Distribution of (a) the normalized number of cherries, (b) the normalized external branch length of the coalescent tree and (c) the time before the MRCA of* 1000 *simulations with no mutation. The means of the distributions are represented by red lines, and the distribution assuming Kingman’s coalescent are represented by dashed red lines.*

Fig E.2 represents the results of the tests presented in Section E.1 depending on the number of simulations, for samples of 100 simulations (as will be done in Section 3.2.2 in the body of the paper). We see that most tests accept neutrality, except Shapiro-Wilkinson test once, but this test is known to be overly conservative when the size of the sample is a bit large.

**Figure E.2:**
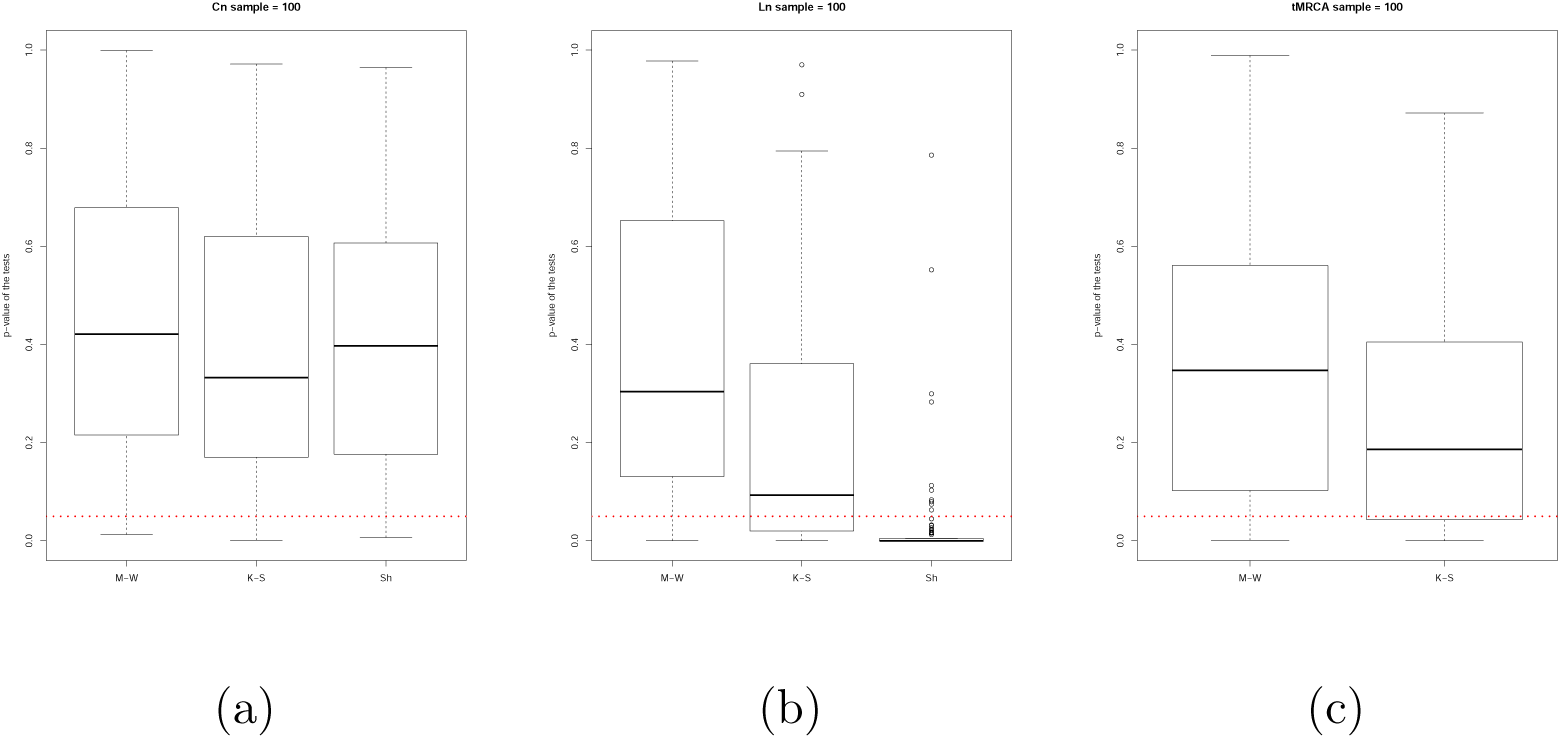
*Tests of neutrality of three statistics depending on the number of sampled simulations: (a) distribution of normalized number of cherries (C*_*n*_*) (b) distribution of normalized external branch length of coalescent trees (L*_*n*_*) and (c) time before the MRCA. Box-plots of the p-values of Mann-Whitney (M-W) and Kolmogorov-Smirnov (K-S) and Shapiro (Sh) tests for* 100 *random samples of* 100 *simulations are shown. The threshold value of rejection of H*_0_ *(0.05) is represented by the dashed red line. If the p-values are inferior to this threshold, the distributions are statistically different from the targeted distribution under H*_0_. *The p-value of the tests computed on all simulations are depicted by the blue symbols.*

### E.3 Neutrality test for the model with mutation

**Figure E.3:**
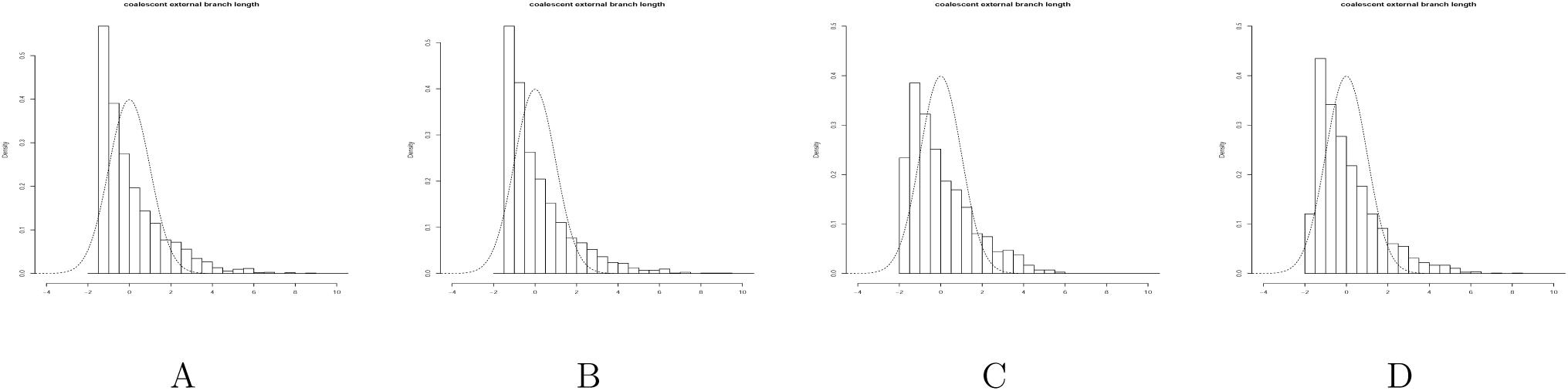
*Histograms of the renormalized external branch lengths produced by the ABC on the pseudo-data A to D. The dashed line represents the distribution followed by Kingman coalescent (Gaussian distribution)*

**Figure E.4:**
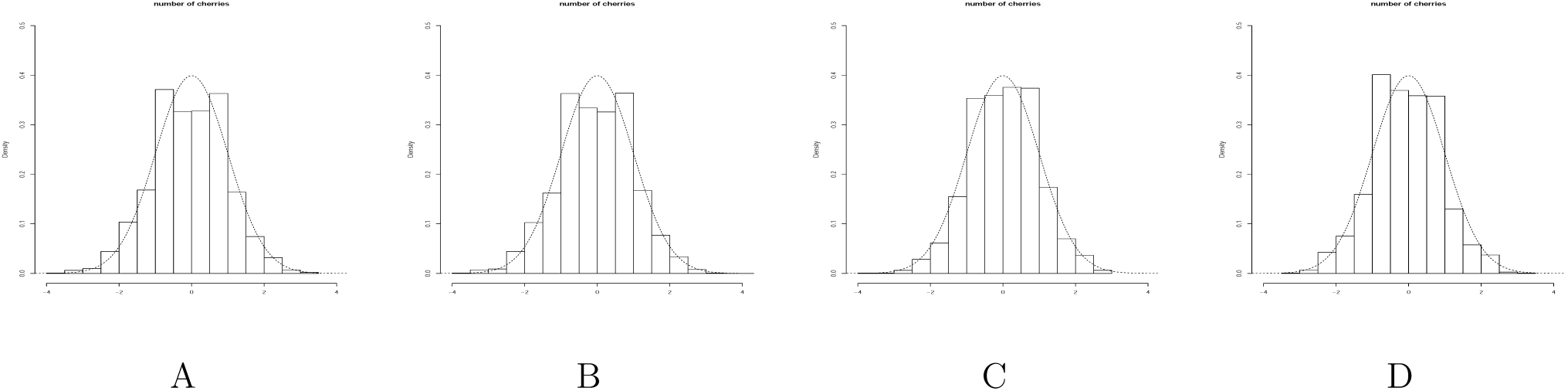
*Histograms of the renormalized number of cherries produced by the ABC on the pseudo-data A to D. The dashed line represents the distribution followed by Kingman coalescent (Gaussian distribution)*

**Figure E.5:**
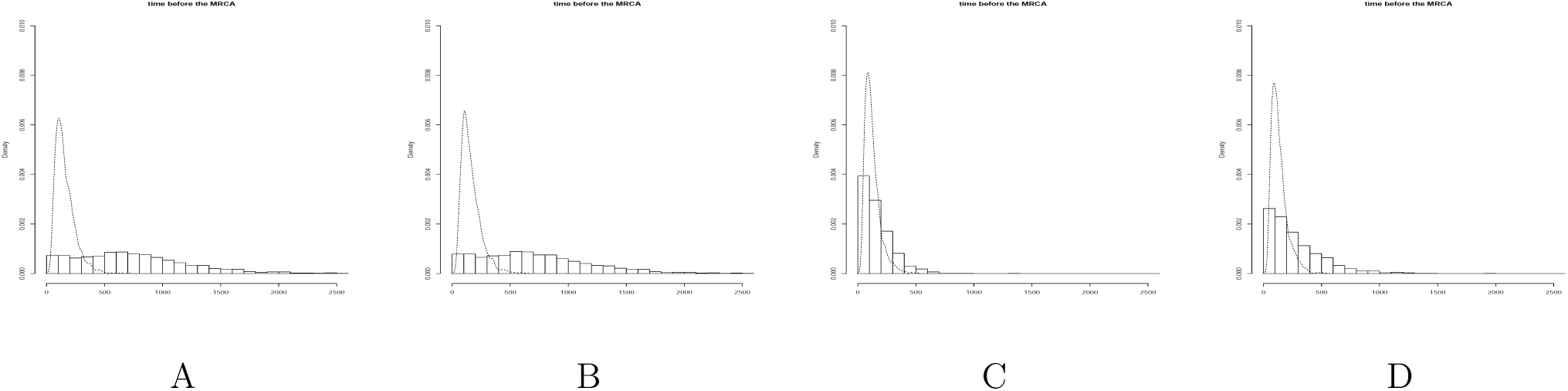
*Histograms of the time to the MRCA produced by the ABC on the pseudo-data A to D. The dashed line represents the distribution followed by Kingman coalescent (obtained by simulations)*

## F Application to patrilineal and cognatic populations in Central Asia

### F.1 Distances between populations

In Figure 3.5(a), it appears that the populations that are considered are distributed roughly along a curve that is plotted in Fig. 3.5(b). The interpolation curve corresponds to a polynomial of degree 3 giving the latitude *y* as a function of the longitude *x*, and that is fitted by ordinary least squares:

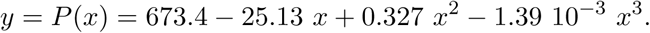

The *R*^2^ associated with this regression is 92.52%.

The populations are then projected on this curve and the distances between two locations are then computed using the line integral. Hence, two populations at locations *z*_0_ = (*x*_0_, *y*_0_) and *z*_1_ = (*x*_1_, *y*_1_) on the graph of *x 1*↦ *P* (*x*) are considered at distance:

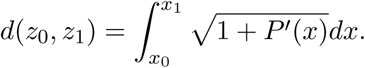

### F.2 Test of the ABC procedure

Our goal is to test the null hypothesis

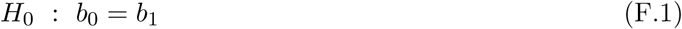

with the alternative hypothesis *H*_*a*_ : *b*_0_ < *b*_1_. For that 20,000 simulations have been performed with parameters randomly drawn in their prior distribution with an additional constraint that either *b*_0_ = *b*_1_ or *b*_0_ < *b*_1_. Two hundred pseudo-data sets have been used to test our ABC procedure. We first used the ABC procedure to estimate the posterior probabilities (see Fig. F.1). Our results show that for most parameters, the estimate parameters are close to their true values.

**Figure F.1:**
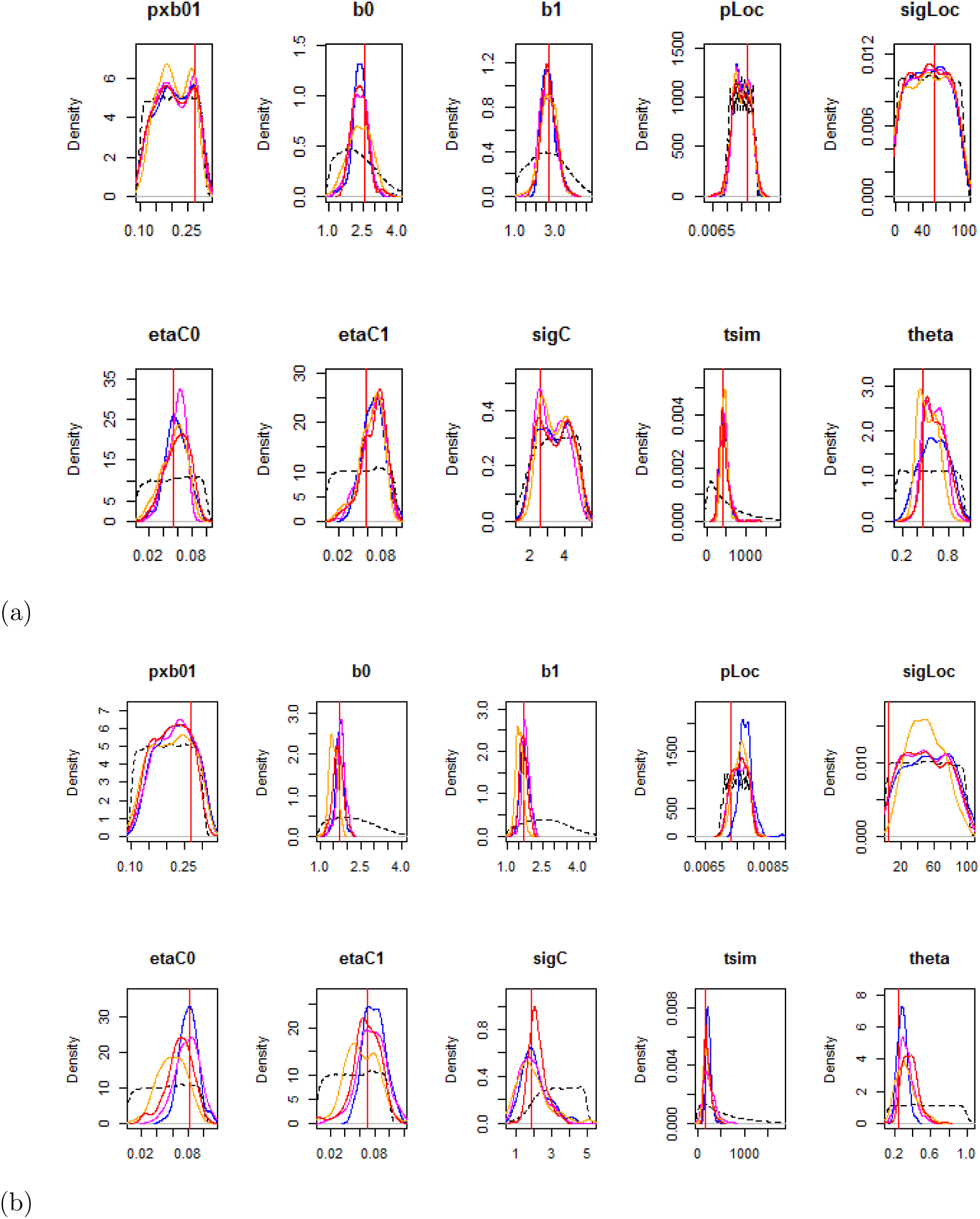
*Results of the ABC estimation for one of the simulated datasets. (a) when the dataset satisfies b*_1_ > *b*_0_, *(b) when the dataset satisfies b*_0_ = *b*_1_. *Legend as in Fig.3.2.*

Second, the approximate posterior probabilities that *b*_0_ = *b*_1_ and *b*_0_ < *b*_1_, respectively noted 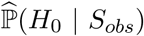 and 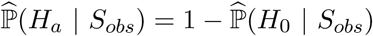, were computed such that

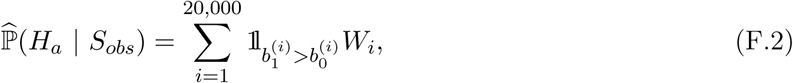

Where 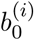 and 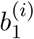 are the birth rate in the simulation *i* ∈ {1, … 20, 000}, and where *W*_*i*_ is the weight defined in App. B. We compute these quantities (F.2) for the real data from Central Asia, but also for 200 ‘training datasets’: 100 datasets chosen from the simulations with *b*_0_ = *b*_1_ presented above and 100 datasets chosen from the simulations with *b*_0_ < *b*_1_. This is represented in Fig. F.2 (a).

The *H*_0_ hypothesis is rejected if 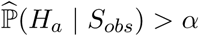 where *α* is a threshold parameter chosen in order to minimize both Type I and Type II errors. For this, we will use the 200 ‘training datasets’ defined above. This allows us to compute the Types I and II errors for different *α* (Fig. F.2(b)). According to these results, we fixed *α* = 0.30 (see caption of Fig. F.2). When *α* = 0.30, Types I and II errors based on the 200 tests performed on simulations are shown in Table 2.

**Table 2:**
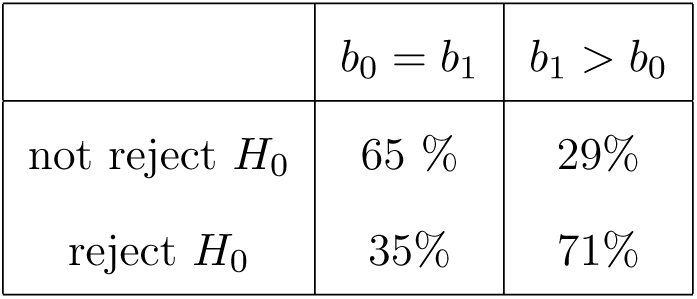
*Confusion matrix based on the 200 tests performed on simulated data with α* = 0.30. *The tests are based on ABC estimation using 20,000 simulations: half of them with the constraint b*_0_ = *b*_1_ *and the other half with b*_1_ > *b*_0_. *Among these simulations, 200 have been used in turn as data: 100 with b*_0_ = *b*_1_ *and 100 with b*_1_ > *b*_0_. *The 200 simulations have been chosen among those where both social models survive with a proportion of at least 10%.*

In Figure F.3, we recover that when *b*_1_ > *b*_0_, the higher the gap between these two values and the more powerful the test is. Graphically, the probability of Type II errors (not rejecting *H*_0_ when *b*_1_ > *b*_0_) is below 5% when *b*_1_ − *b*_0_ > 0.58.

One of the main reason why the test of *H*_0_ is not very powerful is certainly because in most of the simulations, only one social model survives: this happens for 8,862 simulations with *b*_0_ = *b*_1_ out of 10,000 and for 9, 509 simulations where *b*_0_ < *b*_1_ out of 10,000. Although the 200 ‘reference simulations’ are chosen among cases where each social model has a proportion between 10% and 90%, this is hard for the posterior distribution to distinguish differences in fitnesses. This is due to our ecological assumptions: since we supposed that social organization changes occurs in a single direction to state 0 to 1, and since we assumed birth rates such as *b*_0_ ≤ *b*_1_, with a high probability, patrilineal population are expected to go extinct. This makes particularly difficult the use of the ABC procedure in such a case. In addition, for statistical hypothesis testing, the two alternatives are not treated symmetrically, implies that we are lead to choose the threshold *α* appearing in the critical region not necessarily equal to 1/2.

**Figure F.2:**
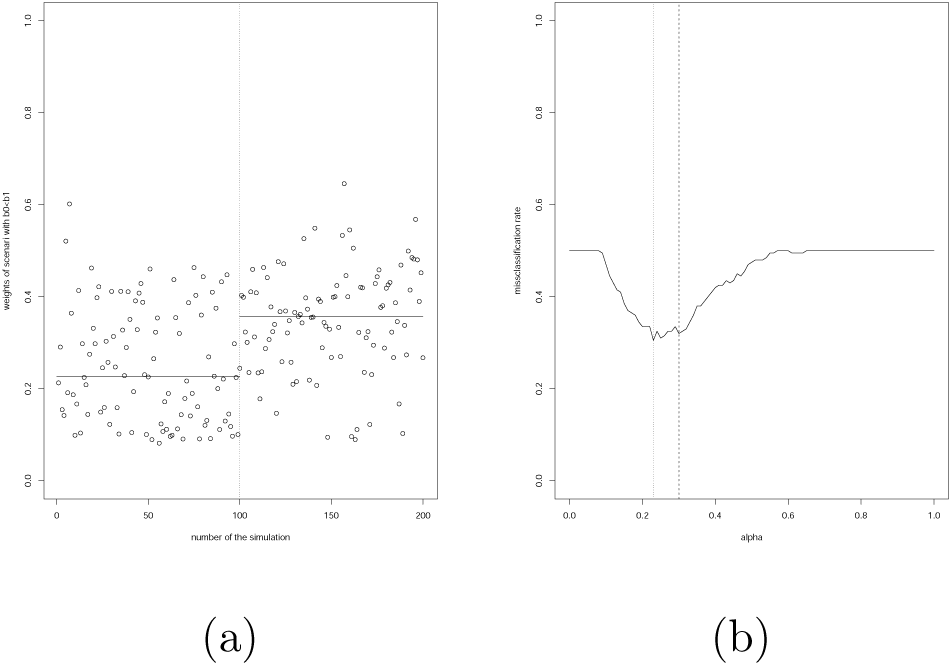
*(a): Estimated posterior probability that b*_0_ < *b*_1_ *conditional on the summary statistics, for each of the 200 simulations chosen in turn to be the ‘data’. The index of the simulation is in abscissa. The 100 first simulations have been obtained under the constraint b*_0_ = *b*_1_ *while the 100 last ones are under the constraint b*_0_ < *b*_1_. *For each simulation, the posterior probability* 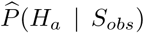 *is computed. The two plain horizontal lines correspond to the medians of these estimations for the 100 simulations where b*_0_ = *b*_1_ *and for the 100 simulations where b*_0_ < *b*_1_: *these medians are respectively* 0.23 *and* 0.36. *(b): The sum of Type I and Type II errors is plotted as a function of the threshold α defining the critical region:*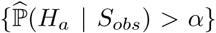. *The Figure (a) entice us to choose the middle of the two medians, α* = 0.3 *(dashed line), as a threshold. The minimum of the sum of the errors on Figure (b) corresponds to α* = 0.23 *(dotted line). Given the precision of the curve close to its minimum, the choice of α* = 0.3 *seems reasonable.*

**Figure F.3:**
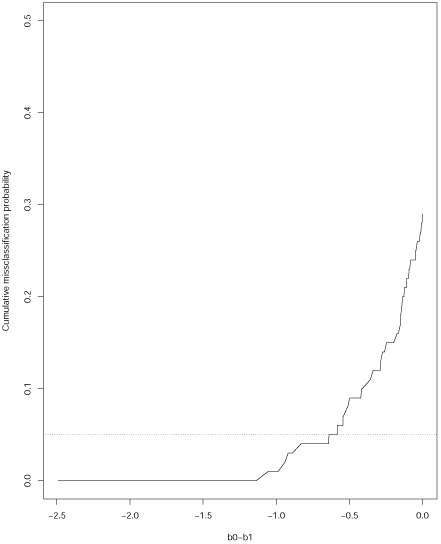
*Cumulated proportion of tests where a Type II error is done, as a function of b*_0_ − *b*_1_, *using the 100 tests performed on simulated data with b*_1_ > *b*_0_.

### F.3 ABC on the Central Asian database

We performed the statistical test (3.1) on the Central Asian dataset. Using the same ABC framework with the same 20,000 simulations, we computed the posterior distribution conditional on the summary statistics. We then performed the ABC test presented as with the pseudo-data sets (App. F.2). Summing the posterior weights of the 10,000 simulations where *b*_1_ > *b*_0_, we obtain for the test statistic:

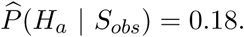

Because this value is much smaller than the threshold *α* = 0.30, we are not in the critical region and *H*_0_ : *b*_0_ = *b*_1_ is not rejected. Hence, there is no statistically significant evidence of fitness difference between the patrilineal and cognatic populations in Central Asia based on genetic data. This can also be seen by comparing the posterior distributions of *b*_0_ and *b*_1_ which look very similar (see Fig. 3.7).

